# The semantics of dreams

**DOI:** 10.1101/2025.05.20.654459

**Authors:** Valentina Elce, Giorgia Bontempi, Serena Scarpelli, Bianca Pedreschi, Pietro Pietrini, Luigi De Gennaro, Michele Bellesi, Giulio Bernardi, Giacomo Handjaras

**Affiliations:** MoMiLab Research Unit, IMT School for Advanced Studies Lucca, Lucca, Italy; Department of Psychology, Sapienza University of Rome, Rome, Italy; School of Biosciences and Veterinary Medicine, University of Camerino, Camerino, Italy

## Abstract

Dreams are universal yet deeply personal experiences. While memory and personal concerns influence dream content, the impact of other individual, generalizable traits remains poorly understood. To address this gap, we built a multimodal dataset including dream and wakefulness reports, alongside demographic, psychometric, cognitive, and sleep-related measures in a large adult cohort. Natural language processing characterized the semantic features that quantitatively distinguish dream from wakefulness reports, with this distinction significantly modulated by individual-specific factors. Longitudinal and cross-sample analyses further demonstrated that major external events, such as the COVID-19 pandemic, affect dream content, leaving lasting traces. Overall, the findings highlight a dynamic interplay between stable individual traits and external events in shaping dream experiences, offering novel insights into the cognitive and emotional architecture of dreaming.

## Introduction

Dreams have long captivated human curiosity, as they are both universal and deeply personal experiences, intimately connected to waking life, yet often strikingly distinct from it. When we sleep, our dreams transport us into immersive worlds - sometimes mundane, sometimes surprising, and occasionally delightful or terrifying. While the artistic, cultural, and psychological significance of dreaming is undeniable, dreams have also gained increasing attention as a subject of rigorous neuroscientific study in recent decades ^1–3^. Indeed, dreams have been proposed not only as a unique model for studying the emergence of consciousness in the human brain but also as a potential window into the functions of sleep itself ^1,4^. As subjective conscious experiences arising during sleep, dreams emerge when the brain is at least partially disengaged from external reality, unfolding as dynamic narratives shaped by prior waking experiences, personal beliefs and expectations, and neural mechanisms that are only beginning to be fully understood ^5,6^.

The analysis of dream contents following experimentally controlled manipulations of individuals’ experiences has led to the hypothesis that dreams, rather than being a mere byproduct of neural activity during sleep, may serve an evolutionary function, related to the mechanisms of learning ^7^, memory consolidation ^8^, or emotion regulation ^9^. According to this view, dreams may aid the individual to cope with waking life events, understand them, and learn from them.

Despite long standing recognition that dreams are shaped by past experiences, their inherent variability and idiosyncrasy have often been seen as obstacles to systematic, comparative analysis. While previous research has shown that common themes may emerge across individuals throughout their lives, individual dreams are still largely viewed as deeply personal, shaped by the dreamer’s unique history, experiences, and concerns ^10^. In this study, we leveraged large-scale, multimodal data and advanced computational linguistic methods to test the hypothesis that not only past experiences but also stable individual traits such as cognitive abilities, personality, and sleep patterns consistently shape the content and structure of dreams. By analyzing thousands of dream and wakefulness reports collected over four years, we investigated how personal traits and collective experiences interact to mold the phenomenology of dreaming, offering a new framework to bridge individual idiosyncrasy with common patterns of dream formation.

## Results

We applied natural language processing tools to analyse two distinct datasets of verbal reports: a multimodal *discovery* dataset, collected from a large sample of adults under typical conditions, and an independent *test* dataset, collected during the COVID-19 lockdown. In both datasets, participants provided daily dream reports over a two-week period via an experience sampling procedure ^11^, with reports collected each morning immediately after awakening. The *discovery* dataset also incorporates reports of wakefulness experiences, daily collected through a further sampling. Participants contributing to this dataset also completed a battery of psychometric questionnaires and cognitive tests (see Materials and Methods), and their sleep-wake patterns were monitored using actigraphy. We used the *discovery* dataset to analyze the semantic content of subjective experiences under physiological conditions while identifying influencing trait and state variables. In contrast, the *test* dataset was used to assess the impact of external large-scale stressors, the COVID-19 pandemic, on dream content. Data collection for the two datasets took place between March 2020 and March 2024 resulting in the inclusion of 287 adult participants (age range: 18–69 years; 110 male participants). In total, 2,038 dream reports and 1,679 wakefulness reports were analyzed.

To capture the semantics of dreams and its relationship with waking experiences, we employed two distinct approaches. The first focused on broad semantic dimensions to identify overarching features of the narratives, while the second examined specific word choices, grouped into lexical domains, to map the detailed content of each report.

### Lexical domains and semantic dimensions

First, we assessed the contribution of semantic dimensions and lexical domains in characterizing verbal reports, and then we explored the interactions and synergies of these intertwining features. In the *discovery* dataset (207 participants; 1,687 dream and 1,679 wakefulness reports), we measured 16 hypothesis-driven semantic dimensions, capturing global narrative features, such as references to different perceptual modalities or spatial details within each text (table S1). The evaluation of each report was performed through three Large Language Models (LLMs), whose performance was validated against four human external raters. We verified the ability of this procedure to capture the subjective semantic nuances of dream experiences by testing the alignment between dimension scorings provided by external observers and the subjective ratings provided by an independent sample of participants who scored their own dreams (see Materials and Methods and Supplementary Figs. S1-S2). Moreover, we identified 32 low-level lexical domains which further clustered into three high-level categories: (i) *Environment*-related domains encompass the visual, geometric, and structural properties of natural and artificial spaces, including colors, objects, spatial locations, and descriptions of buildings and nature. (ii) *Community*-related domains reflect societal structures, institutions, and dynamics, incorporating terms related to education, healthcare, and culture. (iii) *Individual*-related domains capture mental states, emotions, and personal reactions, referencing evaluation processes, fear, and imagination.

In the description of dream experiences (Fig. 1A, lower panel), dimensions reflecting perceptual (i.e., *tactile* and *visual*) and spatial references (*space*) were positively associated with *environment*-related domains (q<0.05). Interestingly, *individual*-related domains exhibited a negative relationship with emotional *valence*, indicating that dreams with stronger internal focus tended to be more negative and with a higher *arousal*. For instance, more distressing and emotionally charged dreams clustered within the *thriller* domain, encompassing themes of danger and life-threatening situations. Further, *auditory* and *communication* references were particularly linked to *community*-related domains, suggesting a connection between verbal interactions and social elements within the reports. Additionally, narratives characterized by heightened *arousal*, frequent *setting* changes and increased *time* references tended to fall within a *drama* domain - an intricate blend of suspense, horror, romance, and tragedy, enriched by storytelling-related terms. Hence, the hierarchical structure of lexical domains aligned closely with the distribution of semantic dimensions, highlighting the intertwining, yet complementary, nature of these features. Each linguistic form, that is the words with their conventionally associated meanings, mapped onto broader semantic aspects, collectively shaping the narratives ^12^.

**Fig. 1.**
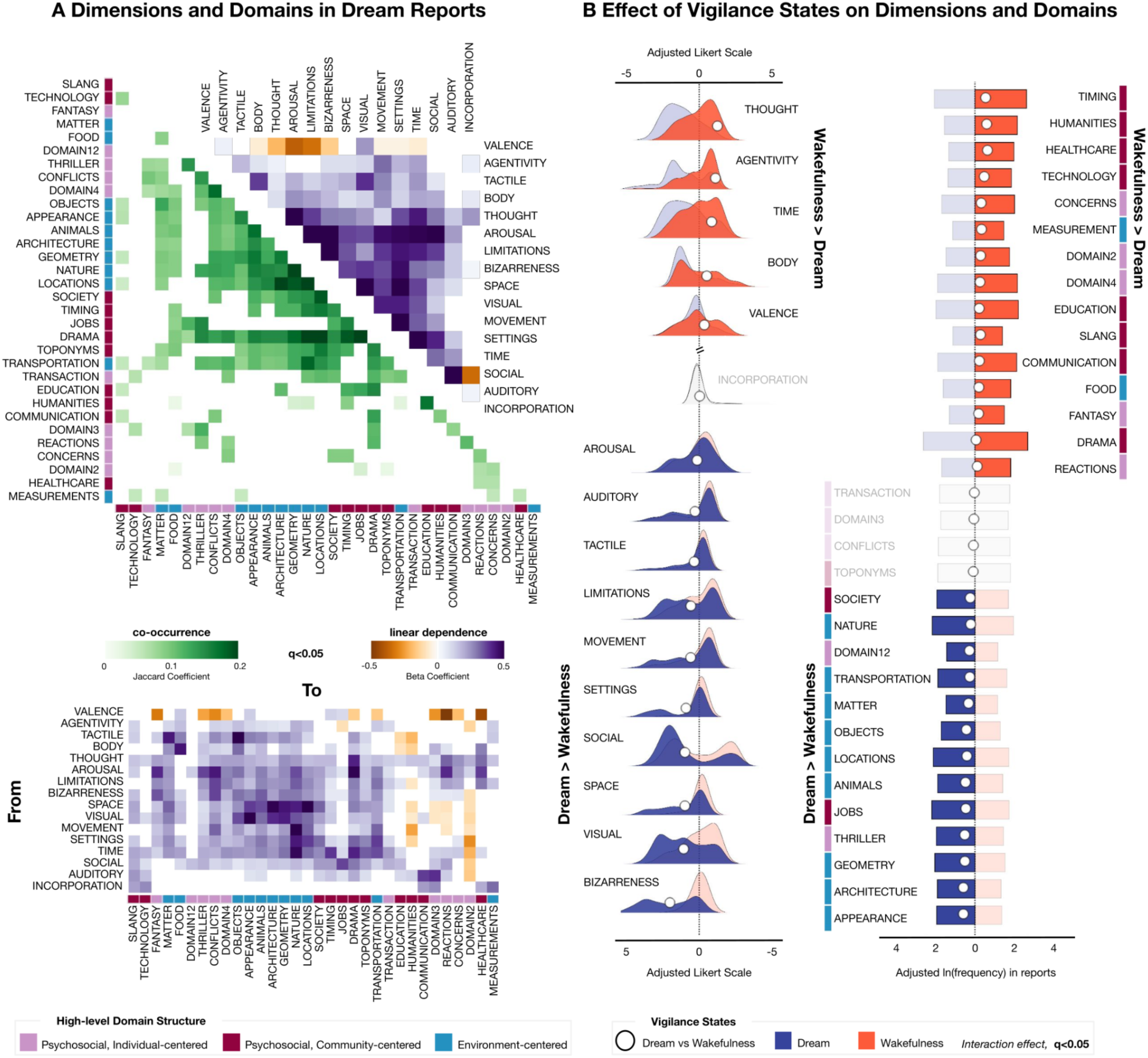
Content analysis of dream and wakefulness reports. Descriptive statistics of report content across vigilance states (i.e., wakefulness and dream). **Panel A**. In the upper section, matrices display associations in dream reports only (through Generalized linear mixed-effect -GLME- model, using age, sex, education level, and verbosity (BADA score) as covariates of no interest; q<0.05, False discovery Rate -FDR- correction) among semantic dimensions and among lexical domains, and their high-level semantic organization. Matrices report the beta coefficient for dimensions and the Jaccard index to quantify the overlap of domains. In the bottom section, the matrix illustrates the associations between semantic dimensions and lexical domains (GLME model; q<0.05). Empty cells indicate non-significant associations. **Panel B.** Semantic feature differences between vigilance states, i.e. wakefulness versus dream reports (GLME model; q<0.05,). The left section of this panel illustrates the distribution of semantic dimensions (rated on a 1-to-9 Likert scale) for wakefulness and dream reports. To facilitate result interpretation, the y-axis is interrupted, with the upper right representing features higher in wakefulness reports and the bottom left indicating those higher in dreams. The right section shows the distribution of lexical domains, quantified by their frequency (in natural logarithm) of occurrence within reports. Reported Likert points and frequencies are adjusted for age, sex, education level, and the BADA score using GLME model. White dots indicate the differences in scores or frequencies between wakefulness and dream reports. Features with a significant interaction effect (q<0.05) are highlighted in red for wakefulness and blue for dreams, while non-significant features are displayed in gray.

### Content analysis of dream and wakefulness reports

Next, we examined potential differences and correspondences across vigilance states, comparing the semantic content of dream and wakefulness experiences (Fig. 1B). Dream reports exhibited significantly more references to perceptual (e.g., the *visual* dimension with a dream minus wakefulness Cohen’s d of 1.52; q<0.05; Supplementary Table S2) and spatial features (e.g., *space* d=1.40, *settings* d=1.29), whereas wakefulness reports were enriched with descriptions of thoughts and metacognitive processes (*thought* d=-1.77). Emotionally, dreams were characterized by higher *arousal* (d=0.21) and more negative *valence* (d=-0.49) compared to wakefulness experiences. Dreams also contained significantly more references to *social* interactions (d=1.40), exhibited a marked increase in *bizarreness* (d=2.85), and featured more references to *limitations* (d=0.82) on the character’s freedom. In contrast, wakefulness reports reflected greater *agentivity* (d=-1.58), with participants portraying themselves as more in control of their actions, aware of the *time* flow (d=-1.20), and attuned to their *body*’s needs (d=-0.73). Notably, the semantic dimension capturing references to the experimental procedure did not differ between dream and wakefulness reports (*incorporation* d=-0.05, 95th confidence intervals: -0.11 0.01, with an average ± standard deviation across all the reports of 1.2±0.87 Likert points). This indicates that the study design did not introduce systematic biases in the sampling of dream and wakefulness experiences.

Dream reports exhibited a significantly higher prevalence of elements related to the *environment* lexical domain, encompassing both animate (e.g., *animals*, d=0.93, present in 18.4% of dream reports; q<0.05; see Supplementary Tables S3 and S4) and inanimate entities (e.g., *nature* d=0.48, 29.8%, *locations* d=0.84, 27.6%, *objects* d=0.40, 13.5%, and their physical *appearance* d=1.13, 19.9%). References to *measurement* and *food* did not follow this trend. In contrast, wakefulness reports were dominated by elements from the *individual* and *community* lexical categories. Exceptions included terms associated with broad descriptive properties (*domain2*, 10.1%), *thriller*-like themes (21.9%), societal structures (*society,* 20.4%), and occupational references (*jobs,* 32.7%), which were more frequently observed in dreams.

### Effect of individual variables on dream and wakefulness report content

We next examined how individual differences influenced the content of reports, focusing on factors that had distinct effects on wakefulness and dream experiences (i.e., interaction between each predictor and the vigilance state, q<0.05; Fig. 2A; Supplementary Tables S5-S6). Attitude toward dreaming emerged as a key predictor, selectively enhancing several semantic dimensions in dream narratives but not in wakefulness, including emotional *arousal* (interaction β=0.05), *bizarreness* (β=0.07), spatial features (*space*, β=0.07), and *visual* perception (β=0.07). Additionally, this trait predicted references to geometric patterns (*geometry*, β=0.09) and navigation in natural environments (*nature*, β=0.07) to a different degree across dream and wakefulness experiences. Notably, subjective sleep quality (β=-0.09), lower vulnerability to cognitive interference (β=-0.03), and a greater propensity for mind-wandering (β=0.27) each contributed to heightened dream *bizarreness*. In particular, proneness to mind-wandering was also associated with more frequent shifts in dream *settings* (β=0.23), which may increase the subjective perception of bizarreness. Lower perceived sleep quality was associated with references to *matter-* (β=0.22) and *appearance-*related (β=0.17) domains (Fig. 2B). Moreover, individuals with higher visuo-spatial memory abilities reported dreams with an increased frequency of *object* references (β=0.09). Younger participants reported *job*- related details more frequently in dreams compared to wakefulness reports (β=-0.03). Additionally, an evening chronotype was linked to a greater focus on *communication*-related content in wakefulness reports compared to dreams (β=0.03).

**Fig. 2.**
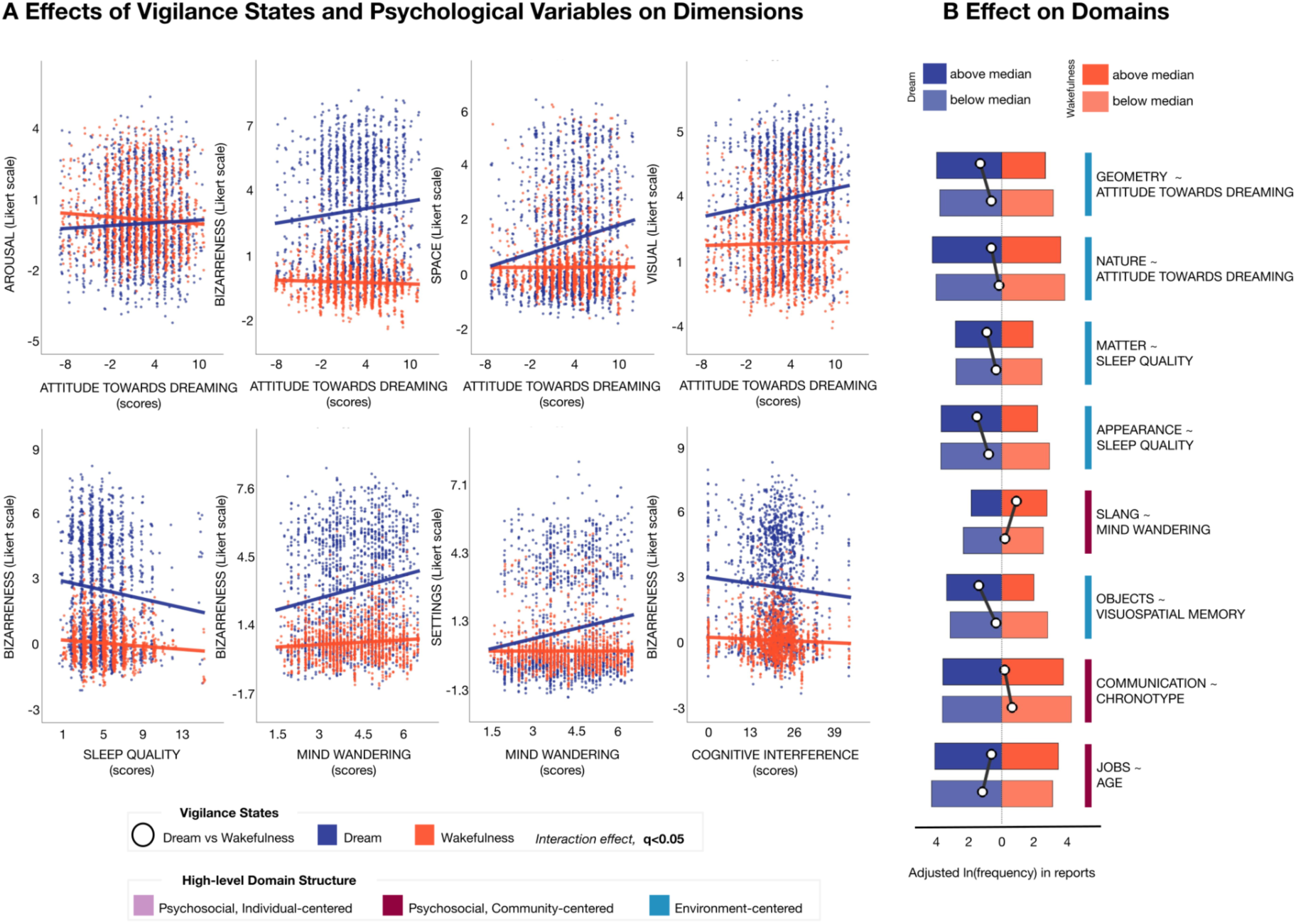
Effect of individual variables on dream and wakefulness report content. Significant effects of individual variables on semantic dimensions and domains (GLME model using psychological variables and their interaction with vigilance states as regressors of interest and age, sex, education level, and the BADA score as covariates; q<0.05, FDR correction). The reported Likert points and frequencies are adjusted for age, sex, education level, BADA score, and all questionnaires and cognitive traits included in the model. Being sleep quality measured through the Pittsburgh Sleep Quality Index, lower scores represent better perceived sleep quality, higher scores worse perceived sleep quality. In **panel A**, the scatter plots show the effect of different predictors on distinct vigilance state dimensions. Each dot represents a different report, with dream reports shown in blue and wakefulness reports displayed in red. Trend lines are also drawn for wakefulness (red) and dream (blue) reports. A slight jittering of points is applied to enhance visualization. **Panel B.** Significant effects of individual variables on lexical domains and their high-level semantic organization. White dots indicate the frequency differences between wakefulness (red) and dream (blue) reports. Bars displayed in darker and lighter colors respectively indicate the reports with the highest (above median) and lowest (below median) values of each predictor.

Further analyses examined the relationship between objective sleep patterns, as derived by continuous actigraphic monitoring, and dream content (Supplementary Fig. S3, Supplementary Tables S7-S8). Among the examined indices, those associated with long, light sleep showed a significant positive association with the frequency of *setting* shifts in dreams (β=0.09, q<0.05).

### Effect of time and external events on sleep conscious experiences

Finally, we examined the impact of the COVID-19 pandemic on dream content using a *test* dataset which was acquired at the end of April 2020 during the lockdown in Italy (q<0.05; Fig. 3A; Supplementary Tables S9-S10). Throughout the period of restrictions, dreams contained significantly more references to *limitations* (d=0.46), *social* interactions (d=0.47), *settings* (d=0.41), *body* (d=0.36), and emotional *arousal* (d=0.50). Additionally, there was an increase in references to fantastical (*fantasy*, d=0.56) or *drama* (d=0.68) elements, work-related themes (*jobs*, d=0.86), and temporal references (*timing*, d=0.53).

**Fig. 3.**
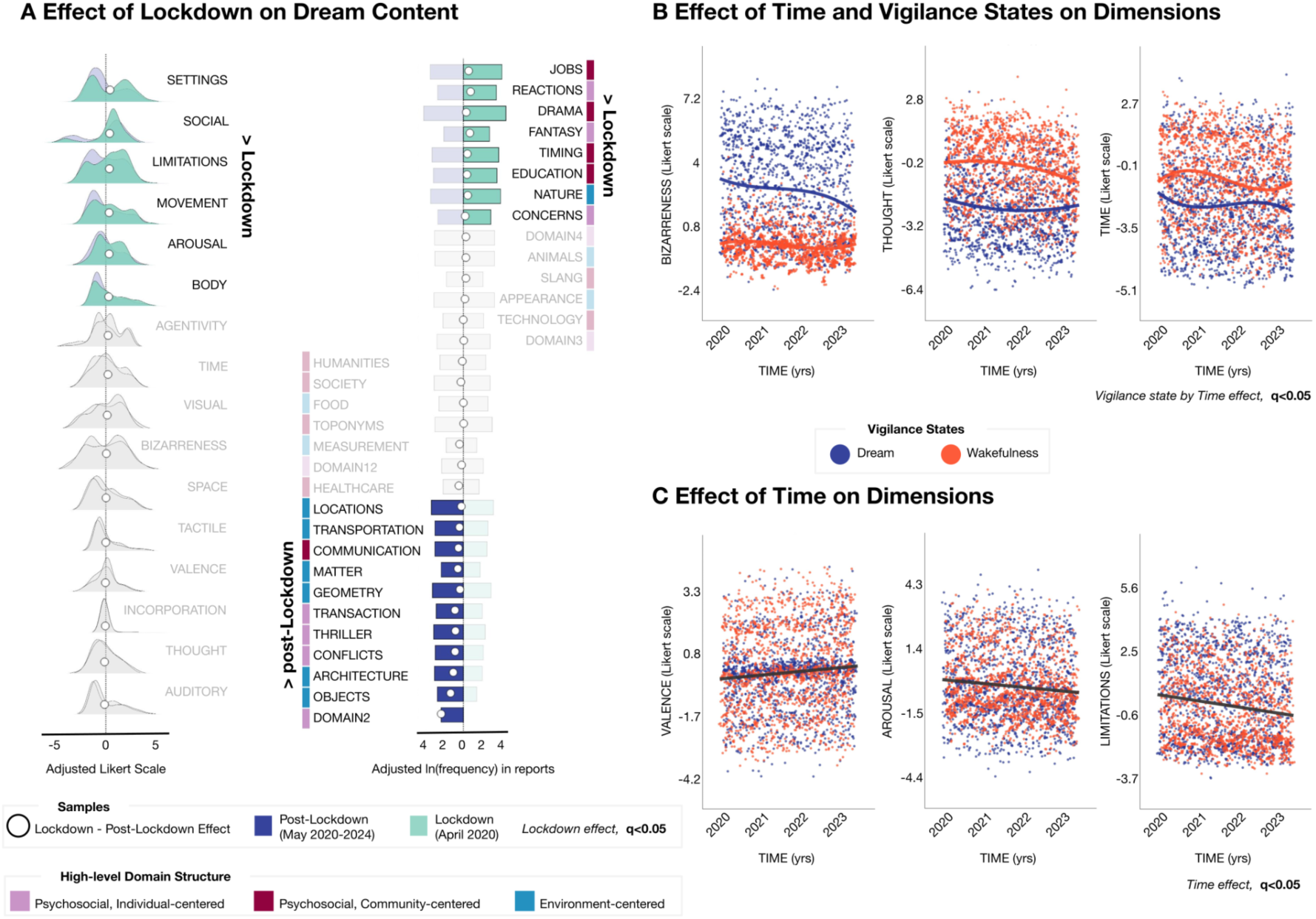
Effect of time and external events on sleep conscious experiences. In **panel A**, comparison between dream reports collected around the time of lockdown release in Italy (test dataset, from the end of April and the beginning of May 2020) and reports of the discovery dataset, collected from the end of May 2020 to March 2024 (GLME model, using age, sex, education level, and the BADA score as covariates; q<0.05, FDR correction). The left column shows the results for semantic dimensions, while the right column shows the results obtained for lexical domains. White dots indicate the differences in adjusted scores or frequencies between lockdown and post-lockdown datasets. Features with a significant effect are highlighted in green for the lockdown dataset and in blue for the post-lockdown dataset, while non-significant features are displayed in gray. **Panel B-C.** Significant effects of the passage of time on semantic dimensions (adjusted scores for age, sex, education level, and the BADA score as covariates using a GLME model; q<0.05, FDR correction). The scatter plots illustrate the semantic dimensions for which we observed an interaction between the passage of time and vigilance states (bizarreness, thought, and time references; **panel B**) or a main effect of the passage of time (limitations, valence, arousal; **panel C**). Each dot represents a different report, with dream reports shown in blue and wakefulness reports displayed in red. Trend lines - obtained by identifying the optimal polynomial fit - are also drawn for wakefulness (red) and dream (blue) reports. On the x-axis, time is represented in normalized units from the beginning of data collection in March 2020 to the end of data collection in March 2024. A slight jittering of points is applied to enhance visualization.

Leveraging the four-year acquisition of the *discovery* dataset, we examined long-term temporal trends in dream and wakefulness reports following the lockdown (q<0.05; Fig. 3B; Supplementary Tables S11-S12). Over time, dream *bizarreness* decreased (interaction β=- 0.74), possibly reflecting a normalization of dream content as pandemic-related stressors subsided. References to *thoughts* (β=0.63) and *time* (β=0.69) also declined, particularly in wakefulness reports, suggesting a shift in cognitive focus during waking life. Furthermore, across both dream and wakefulness reports (main effect of time, q<0.05; Fig. 3C), emotional *valence* tended to increase over time (β=0.59), while *arousal* (β=-0.79) and references to *limitations* (β=-1.09) and to the *society* (β=-0.95) domain progressively diminished.

## Discussion

The generative potential of the dreaming process, its capacity to spontaneously weave vivid, coherent narratives, finds a striking echo in Mary Shelley’s account of the oneiric experience that inspired the Frankenstein novel: “*When I placed my head on my pillow,* […] *my imagination, unbidden, possessed and guided me, gifting the successive images that arose in my mind with a vividness far beyond the usual bounds of reverie*” ^13^. Her words capture the essence of the dreaming mind as both deeply personal and inherently creative. From being the ineffable product of human consciousness, over the past 125 years, dreams have become an object of rigorous scientific observation. The study of how oneiric experiences are shaped by the self and personal experience has progressed from clinical and interpretive approaches focused on individual cases to structured, population-level analyses, where trained raters assess themes, emotions, and characters embedded in dream narratives. These approaches have provided fundamental insights into the nature of dream experiences, highlighting their complexity and connection to waking life. In this study, we leveraged state-of-the-art natural language processing techniques to take this understanding further, demonstrating that dream content is shaped not only by each individual’s unique characteristics but also by broader, generalizable traits and shared external events. This evidence points toward the existence of specific, cross-individual mechanisms by which the brain processes and reconfigures waking experiences during sleep.

Our results show that the overall organization of dream experiences mirrors the narrative structure of identifiable story types found in literature, movies, and plays, shaping a dreamscape that echoes storytelling patterns ^14,15^. Like a movie unfolding on the screen, dreams follow dynamic scene-by-scene structures described at the awakening as vivid, bizarre and immersive. Perceptual details -particularly visuo-spatial elements- dominate the experience, with the dreamer more frequently adopting a spectator-like role, showing a heightened tendency to observe rather than actively engaging in the unfolding events, in contrast to waking experiences. This shift in perspective, combined with frequent setting changes and abrupt conceptual shifts, contributes to the discontinuous, fragmented nature of dream narratives, enhancing their bizarreness in contrast to wakefulness. In line with the view of dreams as *guardians of sleep* ^16,17^, such an immersive and absorbing phenomenological characterization may help explain the heightened sense of disconnection from the external environment during sleep. The vivid and hallucinatory qualities of these endogenous, spontaneously generated contents, indeed, may allow for the suppression of unnecessary sensory inputs by engaging the dreaming cognition with internally generated, rather than externally sourced, information, thus preserving sleep continuity ^18^. Furthermore, while waking experiences appear to be rooted in the daily life dynamics, shaped by the individual’s social interactions, concerns, and insights, dreams tend to be projected in immersive environments. Thus, while wakefulness maintains an egocentric frame of reference - where experiences are anchored to the individual’s physical position and temporal context - dreaming consciousness generates allocentric perceptual simulations. This representational shift effectively decenters the experiential locus, creating immersive scenarios where the dreamer becomes embedded within rather than positioned relative to the simulated environment. Overall, this evidence suggests that waking themes may be transformed during sleep, with elements of reality reshaped and reorganized into largely original oneiric experiences, consistent with the hypothesis that, rather than a direct replay of waking events, dream narratives might offer a hyper associative reinterpretation of lived experiences and future expectations by building broad, non-linear connections among otherwise unexpected or unlikely details ^19^.

Beyond quantifying the phenomenological representation of dreaming, our study revealed how individual trait and state variables shape the content of conscious experiences, exerting distinct effects across vigilance states. While each individual perceives, interprets, and remembers their experiences - whether in wakefulness or in sleep - in a unique way, generalizable traits consistently appear to influence the semantics of dreams. Thus, demographic variables almost exclusively affect the features of verbal reports independently of the vigilance state they describe (see Supplementary Tables S5 and S6), whereas a higher interest in dreams and their significance aids more engaging, immersive experiences during sleep, but not in wakefulness. Interestingly, a positive *attitude towards dreaming* has also been associated with a higher probability of waking up in the morning with at least the perception of having been dreaming just a few seconds before ^20^. Two main non mutually exclusive hypotheses have been proposed to explain the relationship between interest in dreams and dream recall that may be also relevant to explain our present findings ^21,22^. In fact, individuals with a stronger interest in dreaming may present a greater, pervasive focus on their inner experiences and may thus be more likely to remember and report the vivid details of their dreams. It is unclear, though, why this heightened focus would affect only the reporting of specific perceptual aspects of the experience. An alternative interpretation posits that a heightened interest for dreaming might be the surface of a deeper mechanism, yet to be explored, allowing individuals to experience more vivid, immersive dreams. This enhanced dream phenomenology could, in turn, facilitate dream recall and reinforce the individuals’ interest towards dreaming.

Moreover, our findings indicate that dream bizarreness is associated with a higher tendency of the individuals to mind-wander, which also drives frequent shifts in narrative settings. This is in line with accounts suggesting that dreaming and mind-wandering may share a common neural and cognitive foundation. Individuals who frequently mind-wander may have an enhanced propensity to engage in spontaneous, self-generated experiences, independently of external stimuli. Following this reasoning, dreams may represent an intensified form of mind-wandering occurring during sleep. Consistently, we showed in a recent study that proneness to mind-wander is associated with a higher dream recall, potentially reflecting a heightened dream generation process ^20^. While the precise physiological mechanism underlying this increased tendency to engage in internally driven thoughts is yet to be fully understood, some evidence points to a possible role of the so-called default mode network (DMN), a set of brain areas associated with self-reflection and inward thinking ^23^. Our results provide further support to the continuity between waking- and sleep-related mentation demonstrating that a stronger tendency to disengage from external stimuli and focus on the spontaneous flow of internally generated thoughts contributes to shaping the fragmented and discontinuous nature of dreams.

Surprisingly, overall sleep patterns - whether measured objectively through actigraphy or subjectively reported - had a relatively limited impact on dream content. However, specific aspects of sleep did influence particular dream characteristics. Actigraphy-based measures of sleep macrostructure revealed that prolonged light sleep was associated with more frequent shifts in dream settings. Similarly, self-reported sleep quality was related to the bizarreness of dream reports, further supporting the idea that setting shifts contribute to dream bizarreness. These findings align with previous research suggesting that dreams become progressively richer across the night ^4,24,25^. Additionally, lighter sleep stages have consistently been associated with more complex dream content compared to deeper stages ^26,27^. Therefore, longer sleep duration may increase the likelihood of collecting reports of more elaborate and complex dreams. Notably, lower self-reported sleep quality also appeared to influence waking reports more than dream reports in certain aspects. Specifically, individuals who rated their sleep as poorer tended to include fewer references to the appearance and content of their surrounding environment. This observation is in line with evidence indicating that sleep deprivation and poor sleep quality can lead to social withdrawal and a diminished interest in external interactions ^28^.

Our study finally shows how external emotionally salient events, in this case the COVID- 19 pandemic, might affect dream experiences. Dreams are thought to play a crucial role in learning, memory consolidation, and emotion regulation. According to this view, dreaming serves as a mechanism through which the brain processes and integrates newly acquired memories, gradually stripping away or reducing their emotional intensity ^9^. In this light, large-scale stressors -such as the COVID-19 pandemic, which profoundly disrupted daily life on a global scale- could be expected to leave a significant imprint on dream content. Here we found that while narratives during the pandemic retained the immersive and hyper-associative characterization of ordinary dreams, they were more anchored into daily life dynamics and incorporated personal reflections typical of waking experiences. Themes directly related to the pandemic - such as those concerning healthcare scenarios or social interactions - showed no significant changes. However, in a continuous line with what was happening in the daylight world, the actions of the individuals while they were dreaming were described as limited by physical or metaphorical constraints and the recalled emotional states carried a stronger intensity. This suggests that the pandemic modulated specific phenomenological dimensions, likely leading to changes in the hallucinatory depth and in the sense of immersion experienced by the dreamer. The analysis of longitudinal changes across the four-year following the pandemic’s peak period revealed a progressive normalization of dream features, mirroring the epidemiological resolution of the global epidemic. Both dream and waking narratives demonstrated this restorative process, evolving toward more positive affective tones while showing decreasing pandemic-related thematic influence. This temporal pattern suggests that while significant stressors leave measurable imprints on dream phenomenology, these alterations appear to follow a recovering trajectory - diminishing as the psychological impact of the stressor wanes. Notably, this trend aligns with established findings showing parallel normalization of dream experiences and psychological symptoms following traumatic events29,30.

Together, these results bridge longstanding gaps between phenomenological dream research and cognitive neuroscience, offering testable hypotheses about the mechanisms linking dream content to memory consolidation, emotional regulation, and consciousness during sleep. Future studies may build on this framework to develop clinically relevant applications while further elucidating the neurocognitive architecture of dreaming.

## Materials and Methods

This study employed computational linguistics and natural language processing techniques to quantify the semantic features of verbal reports of wakefulness and dream experiences from two distinct prospective datasets. The *discovery* dataset (March 2020–March 2024) captured reports from the general Italian population to analyze the semantic content under physiological conditions while identifying influencing trait and state variables. The *test* dataset (April–May 2020, during the COVID-19 lockdown) served to validate semantic findings from the *discovery* dataset and assess the impact of external large-scale stressors on dream content. Furthermore, two control experiments were conducted. In the first, four trained human raters manually scored dream content to assess the reliability of automatic techniques for quantifying semantic content. The second was a dream diary study, in which participants recorded and later evaluated their own dream experiences. This study aimed to determine the extent to which external human and artificial raters could accurately reflect individuals’ subjective experiences of their own dreams.

### Quantifying the semantic content across dream and wakefulness reports in the *discovery* dataset

This *discovery* dataset was collected to quantitatively analyze the semantic content of wakefulness and dream reports under physiological conditions while identifying individual trait and state variables that may influence these narratives. Given that data collection took place across a four-year period, we also examined longitudinal changes in dream content.

### Participants

The study was conducted on a sample of 217 Italian native language speakers. Data collection was carried out between March 2020 and March 2024. Participants were drawn from the central and northern regions of Italy and their recruitment was conducted through word-of-mouth and flyer distribution. Inclusion criteria were: age from 18 to 70 years old; having regular sleep/wake patterns with six to eight hours of sleep per night, and no diagnosis of sleep-related, neurological, or psychiatric disorders. Exclusion criteria included taking medications that could have affected sleep patterns; having a recent (last six months) history of alcohol and drug abuse, and women who were pregnant, were planning a pregnancy, or were breastfeeding at the time of the study. Ten participants failed to comply with the experimental protocol and dropped out from the study, leading to a final sample of 207 participants (90 male participants; mean ± std, age 34.9 ± 12.4 yrs, range 18-69 yrs; years of education 16.8 ± 2.8 yrs, range 8-23 yrs).

All study participants completed three distinct phases: (i) an initial screening interview followed by the completion of a series of questionnaires, (ii) an experimental period lasting 15 days, during which sleep patterns and verbal reports of subjective conscious experiences during sleep and wakefulness were recorded, and (iii) a concluding session involving an exhaustive cognitive assessment.

The study was conducted under a protocol approved by the Local Joint Ethical Committee for Research (#11/2020). All volunteers signed a written informed consent form before taking part in the study and retained the faculty to drop from the study at any time. No financial compensation was offered for participation.

### Sample Characterization

All volunteers underwent an anamnestic interview aimed at assessing their general health and adherence to inclusion/exclusion criteria. Biological sex, Age and Education were determined by self-report. Recruited participants were then asked to fill out several questionnaires aimed at investigating their attitude towards dreaming ^31^, trait anxiety levels (*State-Trait Anxiety Inventory, STAI*; ^32^), vividness of visual imagery (*Vividness of Visual Imagery Questionnaire, VVIQ*; ^33^), proneness to mind-wandering (*Mind Wandering - Spontaneous and Deliberate Scale*; *MW* ^34^), sleep quality (*Pittsburgh Sleep Quality Index, PSQI*; ^35^), chronotype (*Morningness-Eveningness Questionnaire*; ^36^). Attitude towards dreaming was assessed using a six-item questionnaire where participants were asked to provide their degree of agreement with six statements regarding the general meaning and significance of dreams on a Likert Scale from zero (‘completely disagree’) to four (‘completely agree’). Three items were positive statements about dreams (e.g. ‘dreams are a good way of learning about my true feelings’) and three were negative (e.g. ‘dreams are random nonsense from the brain’). A global score was computed by subtracting the sum of scores provided to the negative statements from the sum of scores associated with the positive statements. Finally, participants evaluated their daytime sleepiness (*Epworth Sleepiness Scale, ESS*; ^37^) and completed a questionnaire extended and adapted from Schredl and colleagues ^38^ concerning their perceived dream recall frequency and several aspects of the dream experiences of the previous three months, including, among others, vividness, bizarreness, and valence. These last two questionnaires were not included in the analyses described in this study.

At the end of the 15-day period, during the final visit, all participants underwent a neuropsychological assessment lasting one hour and aimed to evaluate their cognitive status and to account for potential age-related differences in cognitive functioning (age range: 18-70 yrs). The battery included: *Mini-Mental State Examination*, for a general screening of participants’ cognitive abilities ^39^; *Stroop Task*, for assessing participants’ vulnerability to cognitive interference ^40^; *Babcock Story Recall Test*, for evaluating participants’ verbal memory ^41^; *Rey–Osterrieth complex figure,* for visuospatial memory ^42^; *Phonemic and Semantic Fluency* for evaluating participants’ semantic memory ^43^; *Token Test* focusing on syntactic abilities in language ^44^; *Wechsler Adult Intelligence Scale* (*WAIS*) *- vocabulary subtest* ^45^, yielding information regarding individuals’ semantic knowledge and verbal production and comprehension abilities. Finally, participants were required to complete a free picture description task, extracted from the *Battery for the analysis of aphasic deficits (Batteria per l’analisi dei deficit afasici, BADA*, ^46^), aimed at quantitatively evaluating participants’ connected speech abilities and verbosity by providing them with two pictures representing complex scenes that they had to describe, giving as many details as possible, without any time limit. To obtain a quantitative measure of participants’ verbosity, we computed the word count for the descriptions of both images, averaged the results, and then computed the logarithm of this mean. The complete administration of all neuropsychological tests required about one hour.

Among these, the following tests were included within the analyses: *Stroop Task (cognitive interference)*, *Babcock Story Recall Test (verbal memory)*, *Rey–Osterrieth complex figure (visuospatial memory),* and the verbosity measure extracted from the *BADA* free description task.

### Recording of verbal dream reports and sleep patterns

Participants were provided with an actigraph and a voice-recorder and were asked to record each morning, upon awakening from sleep, everything that was going through their mind just before they woke up, everything they remembered, every experience or thought they had before awakening. Participants were asked to specifically focus on the very last experience they had before the morning awakening in order to minimize the effect of confounding factors that may interfere between the dream experience and its retrieval. Regardless of whether or not they might remember the content of their sleep experiences, participants were required to provide a report. In the event that they woke up with the perception of having been dreaming but could not recall any feature of the experience (i.e. “*white dream*”, ^47^), they were asked to describe this feeling. Similarly, if they woke up with the perception of not having been dreaming at all before waking up, they were asked to report this. Although participants were explicitly instructed to report only the experiences remembered immediately upon awakening, some occasionally recalled and recorded their dreams later in the day. Since these memories could be influenced and altered by external stimuli and waking experiences, we opted to exclude those data from the current analysis (less than 0.1%). After excluding morning reports without descriptions of contentful experiences, the final sample consisted of 1,687 dream reports (reports per participant: 8.1±3.6, min:1, max:16).

Moreover, during the 15 days of the study, participants wore an actigraph to track sleep-wake patterns (MotionWatch-8, Camtech). A subgroup of 50 volunteers (27 female participants, 23 male participants; age 29.7 ± 5.2 yrs, range 22-44 yrs) also had their sleep-related brain activity recorded through a portable Electroencephalographic (EEG) system (*DREEM*) equipped with five EEG dry electrodes (seven derivations: Fp1-O1, Fp1-O2, Fp1- F7, F8-F7, F7-O1, F8-O2, Fp1-F8), a pulse sensor, and a 3D accelerometer. Eight participants interrupted EEG data collection due to discomfort while sleeping. Therefore, we were able to collect neural activity from 42 participants. EEG data were not included in the analyses described in this study.

### Recording of verbal wakefulness reports

With the aim of measuring semantic content differences between wakefulness and dreams, participants were prompted at pseudo-random times throughout the day, once per day, to record the very last experience that was going through their minds up to 15 minutes before. They received a simple text message containing the word “record” (“*registra*”) as a prompt. This approach minimized reliance on long-term memory by capturing thoughts close to their occurrence ^11^. Overall, after excluding promptings which were not followed by a recording (n=238), we collected 2843 wakefulness reports. In this analysis, to account for the potential impact of recent waking experiences in dreams, we downsampled each participant’s wakefulness reports to match the dream reports. Specifically, we selected wakefulness reports from the experimental days preceding recorded dreams; if unavailable, we randomly selected unassigned reports (if available) as substitutes. The random sampling accounted for about 13% of the selected wakefulness reports. The final sample consisted of 1679 wakefulness reports (reports per participant: 8.1±3.5, min:1, max:16).

### Verbal report management and preprocessing

The recordings of participants’ verbal reports were automatically transcribed by means of the Microsoft 365 Word transcriber (Microsoft Corporation). Afterwards, textual data underwent a three-level preprocessing procedure.

At the first level, experimenters manually verified the correspondence between speech and the transcriptions. When needed, they corrected the recordings and added punctuation marks according to the rules of Italian grammar. Verbal reports were also anonymized in order to protect any sensitive data provided by the participants while describing their experiences: any information that would have allowed to identify the volunteers (i.e., proper nouns of cities, places, people) was replaced by a code (e.g., proper nouns of people were replaced by the code [NAME1], [NAME2], etc.).

Afterwards, a second level preprocessing was performed in order to remove any lexical item that did not directly refer to the experiences. In line with previous studies ^48,49^, textual data were manually pruned of volunteers’ commentary about the night (e.g. «*Tonight I slept badly*»), about the task upon awakening and during the day (e.g. «*I remember very few things*», «*What was I thinking before you sent me the message?*»), about the experience itself (e.g. «*That doesn’t make any sense*», «*I don’t know if this is interesting for you*»); linguistic expressions used to introduce the experience (e.g. «*Goodmorning, tonight I dreamt of…*» or, in the case of wakefulness reports, «*Before receiving the message…*»); and false starts, repetitions and self corrections (e.g. «*Kin… kind of of like a boat*»). We also converted arabic numbers into numerals (e.g., the number “3” was converted into the word «*three*»), corrected misspelled words, and regionalisms and dialectal words (e.g. «*and I asked him ‘how ya doin’?’*») were replaced with the Standard Italian equivalent. Afterwards, UTF-8 encoded textual data were fed into three AI models for semantic feature evaluation, as described in the section below.

Moreover, at third-level preprocessing, grammatical categories or parts-of-speech (POS; i.e., categories including, among others, nouns, verbs, adjectives, adverbs, interjections and conjunctions ^50^) were assigned to each word occurring within the verbal reports by means of the *Stanza* toolkit ^50^. Reports and POS tagged texts were then processed, and statistical analyses were performed using MATLAB (The Mathworks Inc., 2024b) as described below.

### AI scoring of semantic dimensions

Second-level preprocessed textual data were fed into three LLMs: LLaMA 3 ^51^, ChatGPT-4, and ChatGPT-4 Turbo ^52^, by using a chat prompting pipeline ^53,54^. The models performed an evaluation of each verbal report across 16 hypothesis-driven semantic dimensions, used to differentiate reports based on meaning and rated on a 9-point Likert scale ^12^. Semantic dimensions were selected based on previous literature and the specific objectives of this study^10,14^. The models were prompted based on the following definitions (for the specific instructions provided, see Supplementary Table S1):

- *Incorporation of experimental procedures* ^55^, that is reporting - either in wakefulness or dream reports - any reference to the experimental protocol the volunteers were asked to perform when participating in the current study (e.g., dreaming of waking up and recording a dream, pressing the event marker of the actigraph, wearing the portable EEG, talking with one of the experimenters or any other character about the study);
- *Thought*, referring to the presence of abstracts thoughts or reasoning of the narrator;
- *Visual experiences*, that is any explicit reference to the sight, but also to details that can be only perceived through it and actions requiring vision, such as looking for somebody in a crowd or aiming at a target;
- *Auditory experiences*, including any reference to the hearing, to details that could be only perceived with it, such as music, noise or voices, and actions that would imply the hearing;
- *Tactile experiences*, meaning not only explicit references to the touch, but also to actions implying the touch, such as brushing, caressing or grasping something with any part of the body, and references to details which can be perceived only with the touch;
- *Valence*, meaning the degree of pleasantness or unpleasantness of the emotional tone of dream and wakefulness reports, irrespective of the emotional intensity, and ranging from extremely negative (e.g. in the case of reports holding feelings of sadness or anger or despair) to extremely positive (e.g. reports carrying feelings instead of happiness, joy or excitement);
- *Arousal*, that is the emotional intensity or strength of reports, regardless of their valence;
- *Bizarreness*, meaning the degree of illogicality, strangeness or discrepancy ^56^ of the events described within the reports;
- *Social processes*, that is any explicit or implicit reference to social interactions, such as talking or arguing with somebody or participating in an activity with other people;
- *Movement*, meaning actions performed by the narrator or any other character within the report that implied movements of the body (e.g. running, jumping, swimming);
- *Space*, that is any reference to details regarding the environment where the events took place (e.g. descriptions of buildings, rooms or landscapes);
- *Change of settings*, including not only explicit references to the displacement of objects or characters from one setting or location to another but also to sudden changes of setting without the explicit reference to an actual movement;
- *Time*, that is references to the chronological aspects of the events, as well as to details regarding their duration;
- *Body*, meaning references to any body part and bodily function or instinct, such as eating, drinking, sleeping or having sex;
- *Limitations of freedom*, meaning elements that restricted or could restrict characters’ freedom of action within the narratives, both physically (e.g. a blocked road or the feeling of wanting to move but being blocked by an invisible force) and in terms of social or moral norms (e.g. refraining from doing something because deemed inappropriate in that particular situation);
- *Agentivity*, assessing how much the narrator actively performed the actions described within the events rather than passively undergoing them.

We computed the median across the scores of the three AIs, obtaining one value for each report and dimension. We also measured the reliability of AI scores in two control experiments (see Supplementary Figs. S1-S2).

### Lexical Domain Analysis

To quantify the semantic content at the lexical level, we developed a procedure based on a bag-of-words representation. First, we used word embeddings to represent the semantic space of all lexical items across wakefulness and dream reports. This semantic space was then decomposed into a set of domains, from which we derived domain embeddings. Next, in each dataset independently, we extracted lemmas and computed the cosine similarity between word embeddings and domain embeddings to generate a normalized lemma-by-domain matrix. This matrix was used to compute a weighted sum of all lemmas per report across domains, yielding a continuous report-by-domain matrix. Finally, we binarized this matrix using a set of null reports to establish binarization thresholds at specified sensitivity and specificity.

### Identification of the Domains

Given the high variability in semantic content, length, and the lack of clear polarization toward specific topics in verbal reports, rather than extracting topics directly from the documents (as done in region models ^57^), we decomposed the semantic space at lemma-level into a set of domains, each centered around a specific topic.

We first identified, through POS tagging, all the lexical items across verbal reports of the *discovery* dataset. We restricted the selection to pronouns and semantically meaningful words: nouns, verbs, adjectives, and adverbs, hence excluding words tagged as conjunctions, determiners and interjections, that is not entailing a lexical meaning but rather a grammatical function ^50^. To obtain numerical vectors for each lexical item, we relied on pre-trained word-embeddings obtained from Wikipedia (snapshot of Italian Wikipedia from 2018-04-20, 300 dimensions, ^58^). Since this version lacked relevant lemmas, we generated new embeddings by averaging those of semantically similar words ^59^. For instance, the embedding for ‘covid’ (which was absent in our Wikipedia snapshot) was derived by merging the embeddings of *‘coronavirus’*, *‘virus’*, *‘pandemia’* (italian word for ‘pandemic’), ‘*contagio*’ (‘contagion’), ‘*influenza*’ (‘flu’), ‘SARS’, and ‘*malattia*’ (‘disease’). Similarly, we built vectors for ‘dreem’ (the EEG device used in the study), ‘*actigrafo*’ (‘actigraph’), and for the anonymization codes ‘PLACE’, ‘NAME’, and ‘INSTITUTION,’ which encoded sensitive information respectively about locations, individuals, and organizations. To approximate lexical information lost during anonymization, we selected representative words for each category, averaged their embeddings, and assigned these values to the respective categories (see Supplementary Table S13 for the specific words used).

By calculating the cosine similarity between the embeddings of all the identified lemmas, we constructed a similarity matrix that captured the entire semantic space. To derive a compact set of components (i.e., lexical domains), we applied a decomposition technique, namely Non-Negative Matrix Factorization (NNMF) ^60^, to obtain a lower-rank approximation of the semantic similarity matrix. We first set to zero all cells in the similarity matrix with a negative cosine value (∼0.04%). Then, following the methodology proposed by ^61^, we decomposed the similarity matrix using a cross-validation procedure to determine the optimal number of components. Specifically, for each tested component count (ranging from 1 to 70), we randomly zeroed out 10% of the similarity matrix cells, estimated the features matrix *W* and coefficient matrix *H* through NNMF (maximum iterations 500, alternating least squares as factorization algorithm), and computed the root mean square error (RMSE) for the left-out 10% by comparing the reconstructed and original similarity measures. Since NNMF is not exact, the procedure was repeated 50 times to obtain a robust RMSE estimate for each component count. We defined the global minimum of RMSE estimates across all the explored components (n=51). Finally, for each domain, we derived a domain embedding by averaging the embeddings of the top 20 lemmas with the highest *H* coefficients. Four raters (GBe, GH, VE, and one artificial intelligence, GPT-4o) labeled each domain by examining the meaning of the lemmas and identifying common themes or shared characteristics among the words in that domain.

To identify high-level macro categories of the domains, we measured the cosine similarity between domain embeddings and applied classical multidimensional scaling ^62^ to the similarity matrix, reducing the representational space to two dimensions. We then performed k-means clustering (with 1000 replications, ^63^) and used the silhouette criterion ^64^ to determine the optimal number of clusters (n = 3), which we labeled as environment-based, community-based, and individual-based macro-domains.

### Domain normalization and evaluation across verbal reports

Since our set of lexical domains was defined solely based on the semantic relatedness of the lexical items, rather than their occurrence or collocations in the reports, we then computed a weighted sum of all lemma weights for each domain and verbal report.

In detail, we extracted semantically meaningful lemmas and computed the cosine similarity between word embeddings and domain embeddings to generate a lemma-by-domain matrix. To enhance the orthogonality between domains, within each domain separately, we sparsified the lemma-by-domain vector by setting to zero all the cosine similarities below a critical threshold determined using the knee-point heuristic procedure ^65^. The rationale behind this procedure was to identify the transition point in the distribution of cosine similarities where the semantic relatedness between lemmas and each domain shifts to a different regime. The procedure resulted in a lemma-by-domain matrix with approximately 98% sparsity, with 50% of the lemmas in the *discovery* dataset (∼5,000) being associated with at least a domain.

After the calculation of the lemma-by-domain matrix, the cosine similarities within each domain were normalized by dividing them by the sum of the nonzero values. Subsequently, within each lemma, data were scaled by dividing them by the sum of the nonzero values, yielding a normalized lemma-by-domain matrix. The final lemma-by-domain matrix maintained the same original sparsity, with nonzero cells containing scaled values up to 1 for lemmas that appeared exclusively within a single domain (e.g., the word *pizza* retained a value of 1 as it was associated only with the *food* domain, whereas *ciotola* -*bowl*- had a value of ∼0.48 for *food* and ∼0.52 for the *objects* domain).

To construct the report-by-domain matrix, we first extracted the lemmas from each report and summed their corresponding weights using the lemma-by-domain matrix. The resulting domain scores were then normalized by dividing them by the total number of lemmas in the report. Since the final report-by-domain matrix contained continuous values, we devised a procedure to binarize the weights, allowing for the estimation of domain frequencies and co-occurrences. Specifically, for each domain, we designed two distinct null distributions of verbal reports: one based on lemmas inherently specific to a single domain (i.e., words with loadings in the lemma-by-domain matrix only for that topic) and another using lemmas shared across multiple domains. The former was used to estimate the number of true positives (i.e., sensitivity), while the latter was used to assess the number of true negatives (i.e, specificity) during the threshold determination for binarization. In detail, the first null distribution was generated by constructing a null dataset with the same number of reports and words per report as the original *discovery* dataset, but ensuring that each null report contained only one domain-specific lemma. From this dataset, we derived the first null report-by-domain matrix. For the second null distribution, we followed a similar procedure but populated each null report with a set of lemmas that were not specific to a single domain, matching their occurrences in the original dataset. This allowed us to construct a second null report-by-domain matrix. Finally, we binarized the original report-by-domain matrix by determining a threshold for each domain at which sensitivity and specificity were equivalent, ensuring that the overall balanced accuracy exceeded 80%. The procedure identified 32 domains out of 51 with an average accuracy of 91±4.2%.

From the binarized report-by-domain matrix, we estimated the absolute frequencies of domain occurrences across dream and wakefulness reports, along with their standard deviations across individuals (Supplementary Table S3).

### Quantify the relationship between dimensions and domains

To measure the association among semantic dimensions and lexical domains in dream reports, we applied three different Generalized Linear Mixed-Effects (GLME) models. As for the dimensions, we created a GLME model (optimizer Nelder-Mead simplex algorithm) using each pairwise combination of dimensions as predictors and predicted variables. Sex, age, educational level, and the outcome of the BADA test were included as covariates of no interest, while the participant was treated as a random effect (see Supplementary Table S14 for a comprehensive description and naming conventions of all the variables collected). The BADA score was incorporated into all examined models as an indicator of individual verbosity. Given that longer reports are more likely to exhibit greater semantic richness, this measure is expected to correlate with higher scores across lexical domains and dimensions. While our primary focus remains on the semantic content of each report, we consider the BADA score to be an independent and reliable metric for assessing an individual’s verbosity. In Wilkinson’s notation, for each pairwise combination *i j* of the dimensions:

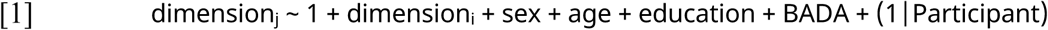

Beta coefficients and p-values related to the dimensions were collected and corrected for multiple comparisons using the False Discovery Rate (FDR, q < 0.05, ^66^). Dimensions were then sorted based on their similarities and presented in fig. 1A, upper section.

A similar procedure was applied to the lexical domains, with the only difference being that we modeled the distribution of the predicted variable as binomial:

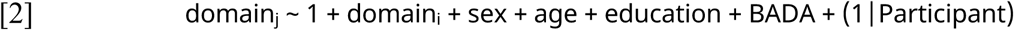

The domains were then sorted based on their Jaccard index, which represented the overlap between each pairwise combination of domains, and models with a significant domain coefficient (q<0.05) presented in fig. 1A, upper section.

Finally, to measure the association between domain and dimensions, we defined with the same approach a GLME model using each domain as the predicted variable, and each dimension as the predictor:

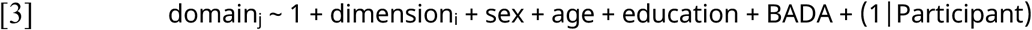

Significant results for the beta coefficient of the dimension (q<0.05) were represented in Fig. 1A, lower section.

### Differences between dream and wakefulness reports

To measure the differences and similarities between wakefulness and dream reports among semantic dimensions and lexical domains, we defined GLME models similar to the previous one, including the vigilance state as a regressor of interest. For each dimension or domain *j*, we defined:

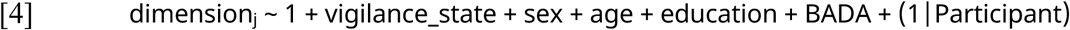

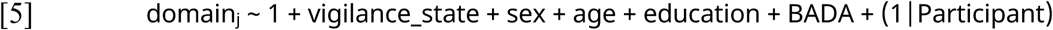

To enhance the robustness of the coefficient estimation associated with the vigilance state, we conducted a permutation test (n=5000). Specifically, we shuffled the vigilance state binary variable within each participant, ensuring that the repeated measures structure of the data was preserved. The non-parametric p-values for the beta coefficient of the report type were estimated by fitting a generalized Pareto distribution to the tails of the null distribution ^67^. Finally, the p-values for the vigilance state coefficients were corrected for multiple comparisons across all dimensions or domains using FDR (q<0.05). The performance measures of the GLME models - including the full-model adjusted R², the full-model parametric p-value, coefficient estimate with the 95% confidence intervals for the vigilance state variable, Cohen’s d, as well as parametric and non-parametric p-value estimates, along with the q-value adjusted for the latter - are reported for each dimension and domain in Supplementary Tables S2 and S4, respectively. The Likert points and frequencies reported in fig. 1B were adjusted for age, sex, education level, and BADA by estimating the residuals - either continuous for dimensions or binomial for domains - using the GLME model.

### Impact of individual characteristics on dream and wakefulness reports

We next examined how individual traits influenced the content of reports, particularly focusing on factors that had distinct effects on wakefulness and dream experiences (i.e., interaction between each predictor and the vigilance state). We defined GLME models that included the vigilance state as the regressor of interest, along with its interaction with the demographic variables mentioned above. Additionally, we incorporated a set of variables derived from questionnaires and cognitive tests. For a detailed description of the collected variables, refer to the Sample Characterization section, Supplementary Table S14 for naming conventions. In Wilkinson’s notation, for each dimension or domain *j*, we defined:

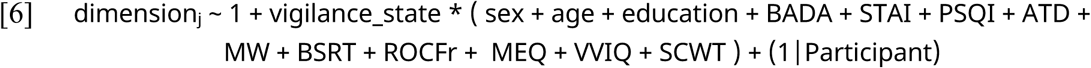

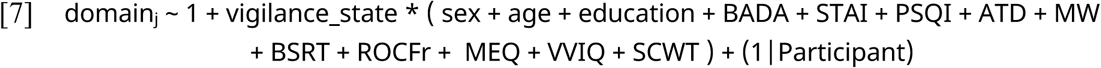

P-values associated with the coefficients were adjusted for multiple comparisons using FDR within each model (q<0.05). The performance measures of the GLME models are reported for each dimension and domain in Supplementary Tables S5 and S6, respectively. The reported Likert points and frequencies in Fig. 2 were adjusted for demographic variables, and all questionnaires and cognitive traits included in the model, excluding the actual predictor used in the plot. Regarding the domains, we represented the interaction effect by dividing wakefulness and dream reports into two subsamples based on the median value of the predictor of interest. Then, we estimated the adjusted frequencies for the associated domain across wakefulness and dream subsamples (Fig. 2B). Results for significant main effects of dimensions and domains are depicted in Supplementary Figs. S3A and S3B, respectively.

### Impact of sleep patterns on dream and wakefulness reports

To investigate the role of sleep macrostructure in shaping dream content, we used actigraphy to obtain objective measures. As detailed in previous work ^20^, we applied Principal Component Analysis (PCA) to reduce 24 actigraphic indices into a set of key components. The indices included, among others, *actual sleep* (or wake) *time* (i.e., the total time spent in sleep/wake according to the epoch-by-epoch wake/sleep categorization) and *sleep efficiency* (i.e., actual sleep time expressed as a percentage of time in bed). This analysis yielded four principal components (PCs), collectively explaining 87.74% of the variance. To aid in interpreting these PCs, we previously used mixed-effect models incorporating sleep structure measures from a subsample of participants who also wore a portable EEG system during the experimental nights. The used indices included the percentages of wakefulness, N1, N2, N3, and REM sleep, as well as the age and sex of participants. Based on the observations and the distribution of PC loadings, the four PCs were labeled as follows: sleep fragmentation (PC1), prolonged non-N3 sleep (PC2; hereinafter referred to as “long, light sleep”), stable sleep with an advanced phase (PC3; “stable advanced sleep”), and unstable sleep with an advanced phase (PC4; “unstable advanced sleep”).

Subsequent analyses focused on dream reports only. We defined GLME models for each dimension or domain by including all predictors (demographics, questionnaires, or cognitive traits) that were found to be significant (uncorrected p-value < 0.05) in the previous analysis, as described above. Additionally, we included the report scores for the first four actigraphic PCs as regressors of interest (see Supplementary Table S14 for naming conventions):

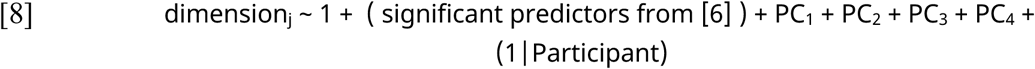

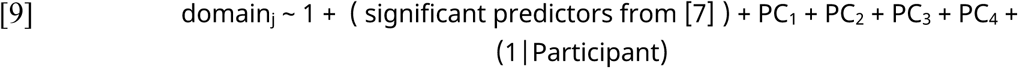

To account for potential collinearities, we included demographics, questionnaires, cognitive traits, and actigraphic data as predictors, as previous research has shown partial interactions among these variables ^20^. P-values associated with the coefficients were adjusted for multiple comparisons using FDR within each model (q<0.05). The performance measures of the GLME models are reported for each dimension and domain in Supplementary Tables S7 and S8, respectively. Results for significant PCs were represented in Supplementary Fig. S3C.

### Impact of time on dream and wakefulness reports

Leveraging the four-year acquisition of the *discovery* dataset, we examined long-term temporal trends in the content of dream and wakefulness reports following the first peak of the COVID-19 pandemic. We defined GLME models similar to [4] and [5], incorporating the vigilance state as a regressor of interest, along with its interaction with time. The variable ‘time’ was coded by sorting all available dates in chronological order and converted into normalized ranks within the 0-1 range. For each dimension or domain *j*, we defined:

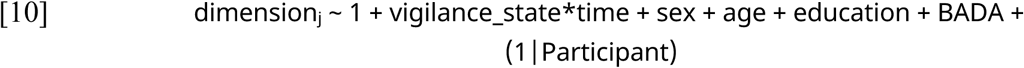

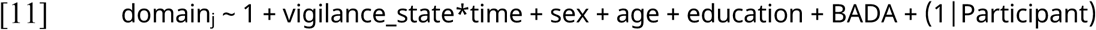

P-values for the interaction between vigilance state and time were adjusted for multiple comparisons across dimensions or domains using FDR (q<0.05). Similarly, the results for the main effect of time were corrected using a separate FDR procedure (q<0.05). The performance measures of the GLME models are reported for each dimension and domain in Supplementary Tables S11 and S12, respectively. The effect of the passage of time on the adjusted variables was represented in Fig. 3B-C. Trends were identified by fitting the optimal polynomial (up to degree 3) to the adjusted data for wakefulness and dream reports independently, when the interaction term was significant (Fig. 3B). For cases with a main time effect, the entire dataset was aggregated prior to trend estimation (Fig. 3C).

### Validating the dream content and quantifying the impact of COVID-19 lockdown on an independent *test* sample

An additional independent (*test*) dataset, gathered during the COVID-19 lockdown in Italy (April-May 2020) ^68^, served two primary purposes: validating the findings related to the overall similarity structure of dimensions and domains, and assessing the potential impact of external, pervasive factors on dream content.

### Participants

During the Italian lockdown, 100 participants were recruited via social media. Only adults residing in Italy with regular sleep/wake patterns and no history of sleep-related, neurological, or psychiatric disorders were included. Those on medications affecting sleep were excluded. Volunteers first completed an online survey collecting socio-demographic data and assessing eligibility. Sex, age, and education were self-reported.

Participants were instructed to keep a dream diary from April 28 to May 11, 2020. The first week (April 28–May 4) corresponded to strict lockdown, while the second (May 5–May 11) marked the easing of restrictions. Of the initial 100 participants, 80 (60 female participants, 20 male participants, age 25.6 ± 4.1 yrs, age range: 19–41 yrs, years of education 14.8 ± 2.1 yrs, range 13-18 yrs) completed the study ^68^.

Similar to the experimental paradigm described above, participants were instructed to record everything that was going through their mind right before waking up everyday, regardless of whether or not they remembered the content of their sleep conscious experiences. Informed consent was obtained, and participants could withdraw at any time. No financial compensation was provided. The study was approved by the Institutional Review Board of the Department of Psychology at Sapienza University of Rome (#0000646/2020).

### Validation of dimensions and domains

From the *test* dataset, we gathered 351 dream reports (reports per participant: 4.4±2.8, min:1, max:14). The reports were treated and preprocessed using the same approach as described for the *discovery* dataset. Regarding dimensions, dream reports were fed into the three LLMs, and ratings were collected following the same procedure outlined above. For the lexical domains, we used the 32 domains identified in the *discovery* dataset and applied the same procedure and parameters described above to generate the weighted report-by-domain matrix. This matrix was then binarized using the thresholds identified in the *discovery* dataset and domain frequencies in reports were evaluated along with their standard deviations across individuals (Supplementary Table S3).

To measure the consistency among dimensions and domains in the *test* dataset, we extracted beta coefficients using GLME models defined in [1] and [2]. Since the BADA score was not available in the *test* dataset, we used the logarithm of the average word count across participants’ reports as a proxy for verbosity. In the *discovery* dataset, this verbosity estimate approximated the original BADA score (Spearman’s ⍴ = 0.168, p = 0.016). Beta coefficients for dimensions and Jaccard indices for domains are reported in Supplementary Fig. S4, along with the similarity between dimensions and domains across the two datasets.

### Impact of lockdown on dream content

Similar to models [4] and [5], we constructed a GLME model that included sex, age, educational level, and the average word count per participant (WC_part) as covariates of no interest, while treating the participant as a random effect. As the predictor of interest, we encoded a binary variable (‘experiment’) to distinguish between reports from the *discovery* dataset and those acquired during lockdown in the *test* set. To assess the impact of lockdown, we compared the *test* dataset with a subsample of the *discovery* dataset, including only reports collected from the end of May 2020. For each dimension or domain *j*, we defined:

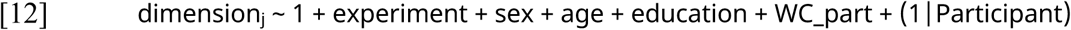

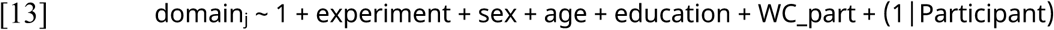

P-values for the experiment coefficients were corrected for multiple comparisons across all dimensions or domains using FDR (q<0.05). The performance measures of the GLME models are reported for each dimension and domain in Supplementary Tables S9 and S10, respectively. The Likert points and frequencies shown in fig. 3A were adjusted for age, sex, education level, and WC_part by estimating the residuals —either continuous for dimensions or binomial for domains— using the GLME model.

### Validating the AI scoring of dimensions

In this initial control experiment, we evaluated the reliability of automatic techniques for quantifying semantic content by comparing computational linguistics measurements with manual dream content scoring performed by four trained human raters.

### Participants and scoring procedure

4 expert human raters (3 females, mean age: 31 ± 5.35 y, all postgraduates with an advanced training in the field of sleep research) were enrolled to score a subsample of the *discovery* dataset including 823 dream reports provided by 86 participants (50 female participants, 36 male participants, mean age: 34.68 ± 11.03), based on the 16 semantic dimensions described above. In particular, the raters were asked, for each report, to answer 16 questions on a Likert scale from 1 to 9. Each question regarded a semantic dimension.

The task was autonomously performed by the scorers on their computers through a custom-made graphical interface that was implemented and presented in MATLAB (The Mathworks Inc., 2022a). Before performing the task, the scorers were provided with written instructions on how to interpret the dimension definitions and answer the questions. The interface automatically saved the scored data at the end of each session.

Each dream report appeared in a white box in the center of the interface. The reports were displayed in random order across the scorers. Below the report, only one question at a time was displayed. Raters scored each semantic dimension by providing their answers using a slider along a continuous scale (Likert from 1 to 9), with the possibility to select values between the displayed labels. Once they answered one question, scorers were asked to press the button “forward” (“*avanti*”) at the bottom of the screen for switching to the next question. At the end of all 16 questions, the following dream report appeared. At the top right of the screen, scorers could also see a counter of the dreams scored within the current session. The median time needed by the scorers to complete the 16 questions for one dream was on average 2’18’’ (range 1’28’’ - 2’54’’), for a total of about 32 hours per rater.

### Measuring the alignment between human observers and AIs

We first assessed the agreement between the dimensional scores provided by the human raters. To account for participant variability, we measured Spearman’s correlation coefficients among raters for each of the 86 participants. For each pairing of raters, we calculated the median of the correlation coefficients across participants to obtain an overall group-level measure. Additionally, we measured the noise-ceiling boundaries by evaluating the correlation of each rater’s scores with the average (median) scores across all four raters to obtain an upper bound, and the correlation of each rater’s scores with the average (median) scores from the other three raters to estimate the lower bound ^69^. Results of the agreement among human raters during the evaluation of the 16 semantic dimensions are shown in Supplementary Fig. S1A.

Similarly to the previous approach, we assessed the agreement between the human raters and each of the three AIs (i.e., LLaMA 3, ChatGPT-4, and ChatGPT-4 Turbo) by estimating Spearman’s ⍴ between the human raters and the AIs for each participant independently. We also estimated the correlation between the median ratings across human raters and the median scores across AIs, with the results reported in Supplementary Fig. S1B.

Overall, these results indicated that the scores provided by the three AIs, particularly when combined, showed a strong similarity with those obtained from external human raters.

### Evaluating the Alignment of Dreamers’ Subjective Ratings with Human and AI observers

The second control experiment was a dream diary study, in which participants recorded their dream experiences for 15 days immediately upon waking and evaluated their own dream content. This study aimed to determine the extent to which external human and artificial raters could reflect individuals’ subjective experience.

### Participants

In this control experiment, 10 Italian native language speakers (6 female participants, 4 male participants, mean ± std, age 30.18 ± 4.7 yrs; range 25-41 y) were recruited. Similar to the paradigm described above, we only recruited individuals with regular sleep/wake patterns, six to eight hours of sleep per night, and no diagnosis of sleep-related disorders or of any other pathological condition that might have compromised their sleep.

### Recording of verbal dream reports, sleep patterns, and subjective ratings

In this study, participants were provided with an actigraph and were required to complete a 15-day dream diary only. Volunteers were provided with the same instructions used in the *discovery* dataset and described above. Here, participants recorded and sent their sleep conscious experience reports to an experimenter via vocal messages.

Immediately after completing the recording, participants were asked to fill out a questionnaire consisting of 23 items divided into two sections. In the first section, if participants were able to recall the content of their dreams upon awakening, they responded to 17 questions on a 9-point Likert scale. These questions assessed the semantic dimensions of their dream experience, as described above, as well as their confidence in the completeness and accuracy of their recollections. The second section of the questionnaire focused on participants’ current mood, level of tiredness, and the perceived quality of their nocturnal sleep, all rated on a 9- point Likert scale. Additionally, they provided information on their falling asleep and awakening times and indicated whether their awakening was spontaneous or induced by external factors such as the alarm, the bed partner or other noises. If participants awoke with the feeling of having dreamt but were unable to recall details of the experience, or if they felt they had not dreamt at all, they were instructed to complete only the second section of the questionnaire, reporting on their mood, tiredness, sleep quality, sleep duration, and the nature of their awakening.

Notably, during the recruitment phase, participants were provided with written instructions regarding the questions they were asked to answer and including the specific definitions of each semantic dimension, which were based on the prompting input to the AI models.

Finally, from a minimum of 30 to a maximum of 40 days after completing the task, participants were contacted again for a follow-up session. They were asked to read the reports of the contentful dream experiences they recorded during the 15-day dream diary protocol and, for each report, to answer the 16 questions regarding the semantic dimensions and assess, on a 9-point Likert Scale, how vivid was their memory of the dream experience. Of note, before the follow-up, reports of sleep conscious experience were treated up to the second-level preprocessing used in the main study and described above. This follow-up task was submitted online and the reports were administered in random order. Overall, in this control experiment we gathered 64 dream reports (reports per participant: 6.4±3.3, min:2, max:12).

### Measuring the alignment between subjective ratings, human and AIs observers

We first assessed the agreement between the dimensional scores provided by dreamers immediately after awakening and after the 30-day follow-up. Similar to the previous approach, Spearman’s correlation coefficient was calculated for each participant, and then the median estimate for each dimension was computed. Results of the agreement among dreamers during the evaluation of the 16 semantic dimensions are shown in Supplementary Fig. S2A.

We then assessed the agreement between dreamers and each of the three AIs (i.e., LLaMA 3, ChatGPT-4, and ChatGPT-4 Turbo) and their combination, using the same approach used above. Results are reported in Supplementary Fig. S2B.

Overall, these results indicated that the scores provided by the three AIs, particularly when combined, showed a strong similarity with the subjective ratings reported by the dreamers.

## Author Contributions

Conceptualization: G.Be., M.B., G.H., V.E.; Investigation: V.E., G.Bo., B.P., S.S.; Methodology: G.H., G.Be., V.E.; Software: G.H., G.Be., V.E.; Formal Analysis: V.E., G.H.; Visualization: V.E., G.Be., G.H.; Data Curation: V.E., S.S., G.Bo.; Validation: V.E., G.Be., G.H.; Supervision: G.H., G.Be., P.P., L.D.; Funding acquisition: G.Be., M.B.; Project administration: G.Be.; Resources: G.Be., L.D., P.P; Writing - Original Draft: V.E., G.Be., G.H.; Writing - Review and Editing: All authors. All authors have read and agreed to the published version of the manuscript.

### Acknowledgements

The authors thank Alessandro Lenci, Davide Bottari and Claudia Picard-Deland for their feedbacks on a preliminary version of the work and for providing constructive comments and suggestions which helped to improve the manuscript, Francesco Lomi, Luca Fuligni, Elena Capriglia, Monica Di Giuliano, Margherita Bozzoli, Federico Frau, Aurora Salina, Damiana Bergamo and Giulia Avvenuti for their help in data collection and preprocessing, and all volunteers for participating in this study. This work was supported by a grant from the BIAL Foundation (#091/2020) and from the European Union - Next Generation EU, Missione 4 Componente 2 Inv. 1.1 CUP D53D23009580006, project PRIN 2022 The Language Of Dreams: the relationship between sleep mentation, neurophysiology, and neurological disorders (2022BNE97C) and the *TweakDreams* ERC Starting Grant (#948891) (to GBe). The funders had no role in the conceptualization, design, data collection, analysis, decision to publish, or preparation of the manuscript.

## Notes

### Competing Interest Statement

The authors have declared no competing interest.

