## Supplementary Materials for "The semantics of dreams"

### Supplementary Figures

Figure S1

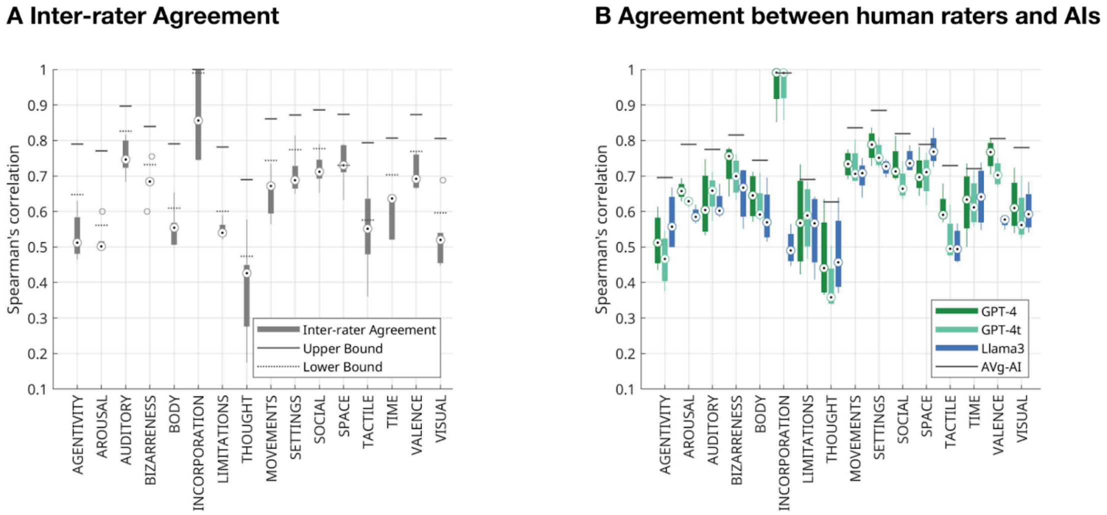


***Fig. S1. Agreement among human raters and AIs in the evaluation of the 16 semantic dimensions.*** *(A) The boxplots represent the Spearman correlation coefficients among raters for each semantic dimension. The noise-ceiling boundaries are also displayed. The upper bound was computed by evaluating the correlation of each rater's scores with the average (median) scores across all four raters. The lower bound was estimated as the correlation of each rater's scores with the average (median) scores from the other three raters. The average agreement was* $\rho$*=0.626±0.115, with a minimum* $\rho$*=0.427 for thoughts, and a maximum of* $\rho$*=0.856 for incorporation. (**B**) The boxplots represent the Spearman's correlation coefficient between the human raters and each of the three AIs (i.e., LLaMA 3, ChatGPT-4, and ChatGPT-4 Turbo) for each semantic dimension. The correlation between the median ratings across human raters and the median scores across AIs is also displayed with a black continuous line with an average agreement of* $\rho$*=0.781±0.085 (minimum* $\rho$*=0.627 for thoughts, maximum of* $\rho$*=0.990 for incorporation).*

Figure S2

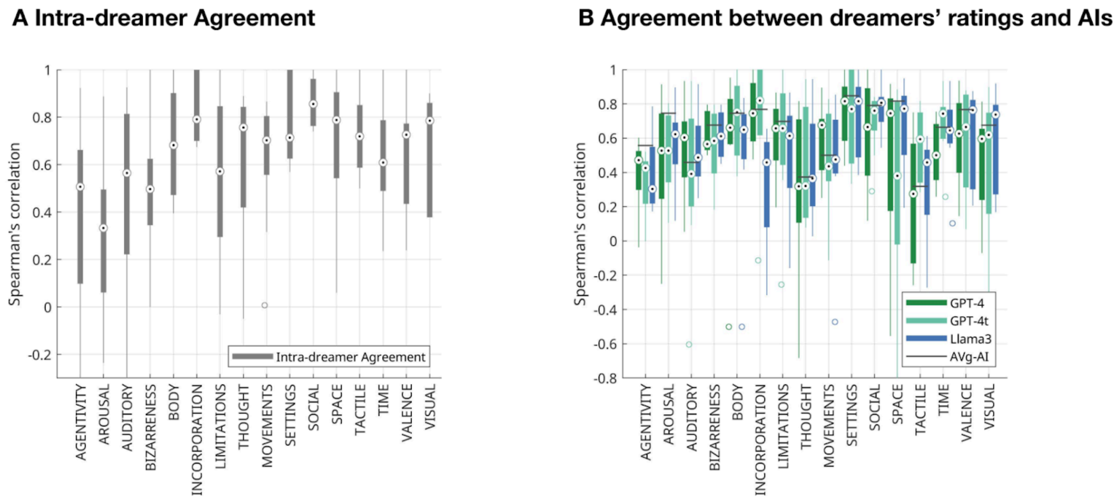


***Fig. S2. Agreement between dreamers and among dreamers and AIs in the evaluation of the 16 semantic dimensions.*** *(**A**) Spearman's correlation coefficient between the scores provided by dreamers immediately after awakening and after the 30-day follow-up for each semantic dimension. The average agreement was* $\rho$*=0.663±0.138, with a minimum* $\rho$*=0.333 for arousal, and a maximum of* $\rho$*=0.856 for social. (**B**) Spearman's correlation coefficient between dreamers and each of the three AIs (i.e., LLaMA 3, ChatGPT-4, and ChatGPT-4 Turbo) for each semantic dimension. The correlation coefficient between the median ratings across dreamers and the median scores across AIs is also displayed with a black continuous line with an agreement of* $\rho$*=0.650±0.161 (minimum* $\rho$*=0.319 for tactile, maximum of* $\rho$*=0.847 for settings).*

Figure S3

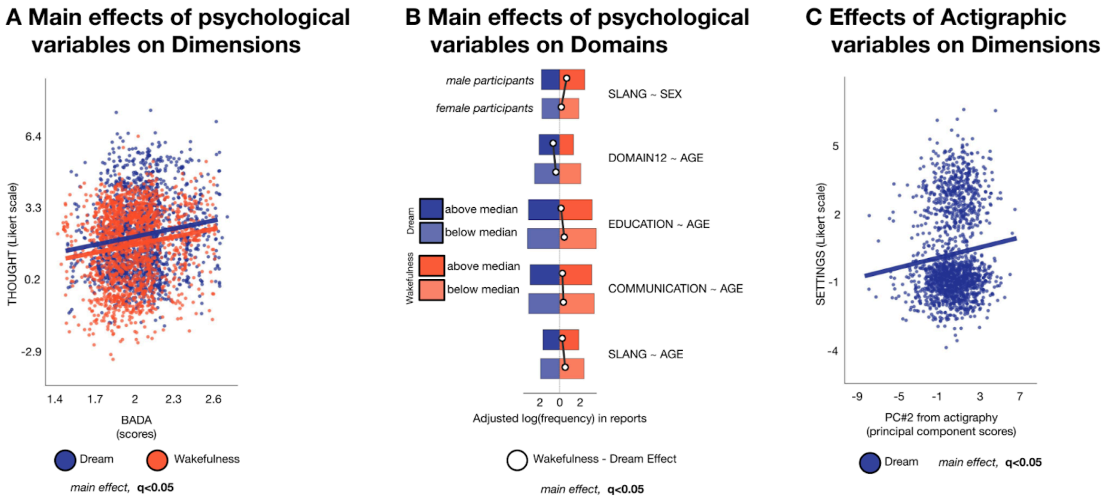


***Fig. S3. Effect of individual variables on dream and wakefulness report content.*** *Significant effects of individual variables on semantic dimensions and domains (GLME model using psychological variables and their interaction with vigilance states as regressors of interest and age, sex, education level, and the BADA score as covariates; q<0.05, FDR correction). (**A**) The scatter plot shows the significant effect of individual verbosity (BADA scores) on the description of thoughts and metacognitive processes (thought dimension) in distinct vigilance states. Each dot represents a different report, with dream reports shown in blue and wakefulness reports displayed in red. Trend lines are also drawn for wakefulness (red) and dream (blue) reports. A slight jittering of points is applied to enhance visualization. (**B**) The bar plot shows significant effects of individual variables on lexical domains. White dots indicate the frequency differences between wakefulness (red) and dream (blue) reports. Bars displayed in darker and lighter colors respectively indicate the reports with the highest (above median) and lowest (below median) values of each predictor. The plot evidences an increased use of slang in wakefulness reports provided by younger male participants, whereas older volunteers showed lower lexical references to education, communication dynamics and domain12. (**C**) The scatterplot shows the significant effect of actigraphy-based PC2 (long light sleep) on setting shifts in dream reports (blue dots); the trend line is also drawn in blue. A slight jittering of points is applied to enhance visualization.*

Figure S4

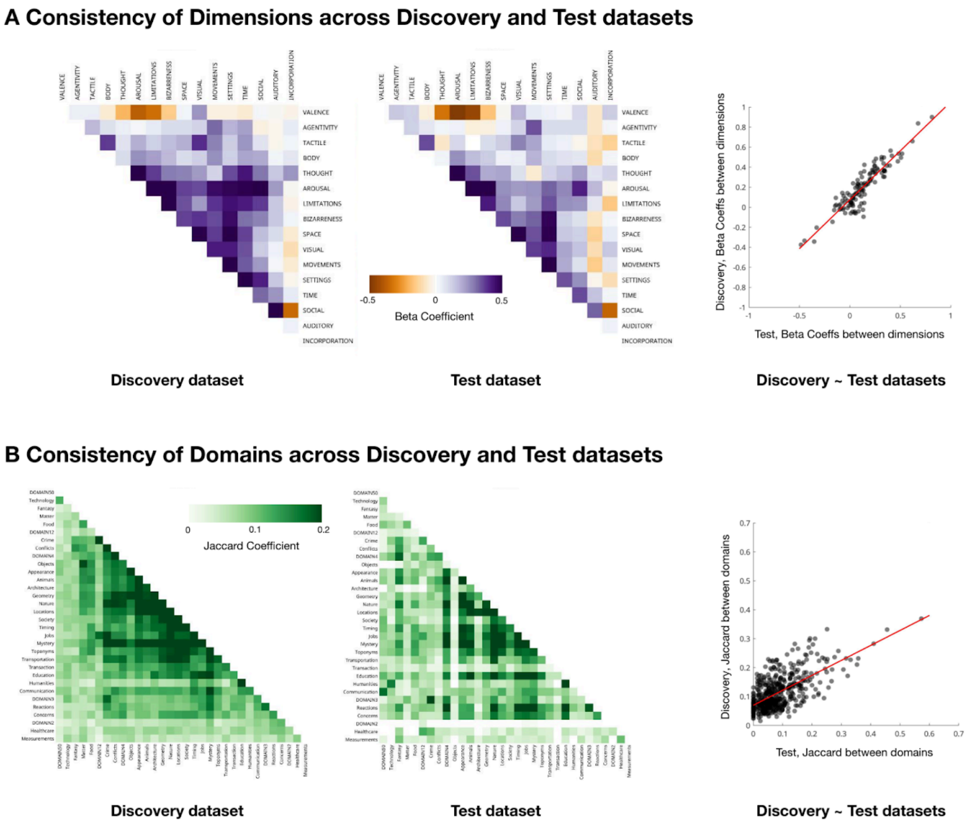


***Fig. S4. Consistency of semantic dimensions and lexical domains across discovery and tests datasets.*** *Beta coefficients for dimensions and Jaccard indices for domains are reported (computed through Generalized linear mixed-effect -GLME- model, using age, sex, education level, and verbosity (BADA score) as covariates of no interest; q<0.05, False discovery Rate -FDR- correction), along with the similarity between dimensions and domains across the two datasets. (**A**) Matrices display the beta coefficient in dream reports only among semantic dimensions in discovery (left) and test (right) datasets. The scatter plot shows the beta coefficients in the discovery and test dataset. The agreement between dimension across the two dataset was* $\rho$*=0.873, p<0.00001. (**B**) Matrices report the Jaccard index to quantify the overlap of lexical domains in discovery (left) and test (right) datasets. Empty cells indicate non-significant associations. The scatter plot shows the Jaccard indices in the discovery and test dataset. The scatter plot shows the beta coefficients in the discovery and test dataset. The agreement between domains across the two dataset was* $\rho$*=0.554, p<0.00001.*

#### Supplementary Tables

Table S1

| DIMENSIONS | PROMPTING |
|---|---|
| AGENTIVITY | "Evaluate how much the narrator of an Italian text is the agent of the actions described on a scale from 1 to 9. The scoring does not concern whether the valence of the actions is positive or negative. With score 1 you need to score text where the narrator is extremely passive and undergoes the actions performed by others. With score 5 you need to score text where the narrator is neither passive or active or does not take part into the actions described. With score 9 you need to score text where the narrator is extremely active and perform the actions. Answer only with a number. Here is the text in Italian: " |
| AROUSAL | "Evaluate the emotional intensity of an Italian text on a scale from 1 to 9. The scoring is not related to the emotional valence. The score only concerns the emotional strength of the narration. With score 1 you need to score text not at all emotionally intense. With score 5 you need to score text moderately intense. With score 9 you need to score text extremely intense. Answer only with a number. Here is the text in Italian: " |
| AUDITORY | "Evaluate how much an Italian text describes auditory experiences on a scale from 1 to 9. You should evaluate how much the situations include talking, hearing, listening and any details that can only be perceived either by the narrator or other characters through the hearing, such as voices, noise and music. With score 1 you need to score text without any semantic relatedness to auditory experiences. With score 5 you need to score text with a moderate semantic relatedness to auditory experiences. With score 9 you need to score text with extreme semantic relatedness to auditory experiences. Answer only with a number. Here is the text in Italian: " |
| BIZARRENESS | "Evaluate the bizarreness of an Italian text on a scale from 1 to 9. You should evaluate how much the situations described in the text are strange or unnatural according to your specific experience. With score 1 you need to score text describing completely normal and common situations. With score 5 you need to score text describing partially strange or unnatural situations. With score 9 you need to score text describing extremely strange or unnatural situations. Answer only with a number. Here is the text in Italian: " |
| BODY | "Evaluate how much an Italian text refers to the body and the bodily functions on a scale from 1 to 9. You should evaluate how much the text includes any kind of reference to body parts and to physiological functions or instincts, such as sleeping, eating, having sex. With score 1 you need to score text without any semantic relatedness to the body or the bodily functions. With score 5 you need to score text with a moderate semantic relatedness to the body or the bodily functions. With score 9 you need to score text with an extreme semantic relatedness to the body or the bodily functions. Answer only with a number. Here is the text in Italian: " |
| INCORPORATION | "Evaluate how much an Italian text relates to any of the following concepts: 'dreams', 'dreaming', 'sleep', 'experiment', 'EEG', 'electroencephalography', 'actigraph', 'actigraphy', 'accelerometer', 'smartwatch', 'recording', 'recorder', 'voice recording'. Provide a score on a scale from 1 to 9. With score 1 you need to score text without any semantic relatedness to these concepts. With score 5 you need to score text with a moderate semantic relatedness to these concepts. With score 9 you need to score text with extreme relatedness to these concepts. Answer only with a number. Here is the text in Italian: " |
| LIMITATIONS | "Evaluate how much an Italian text includes elements limiting or which might limit the freedom of the characters on a scale from 1 to 9. You should evaluate how much the text describes or refers to elements limiting or which might limit the freedom of the characters either physically or in terms of ethical and moral boundaries. The score concerns the presence of details which might represent an obstacle or a limitation for the characters, such as a blocked road, the feeling of not being able to move, or refraining from doing something because it is not allowed or appropriate in a situation. With score 1 you need to score text without any limitations to the freedom of the characters. With score 5 you need to score text with a moderate number of limitations to the freedom of the characters. With score 9 you need to score text with several limitations to the freedom of the characters. Answer only with a number. Here is the text in Italian: " |
| THOUGHT | "Evaluate how much an Italian text describes abstracts thoughts or reasoning of the narrator on a scale from 1 to 9. With score 1 you need to score text without any abstract thoughts or reasonings. With score 5 you need to score text with an average number of thoughts or reasonings. With score 9 you need to score text with several descriptions of thoughts or reasonings. Answer only with a number. Here is the text in Italian: " |

| | |
|---|---|
| MOVEMENT | "Evaluate how much an Italian text describes physical movements on a scale from 1 to 9. You should evaluate how much the situations include any kind of physical movement performed by the characters, including the narrator. The scoring concerns movements which can be only performed with the body, such as running, swimming, flying, jumping. With score 1 you need to score text without any semantic relatedness to physical movements. With score 5 you need to score text with a moderate semantic relatedness to physical movements. With score 9 you need to score text with an extreme semantic relatedness to physical movements. Answer only with a number. Here is the text in Italian: " |
| SETTINGS | "Evaluate how much an Italian text includes changes of the surrounding environment and of the scenery where the events take place on a scale from 1 to 9. You should evaluate how much the text includes moving or displacement of objects and people from one environment or place to another. The scoring also concerns sudden changes from one scenery to another without any explicit description of the displacement. With score 1 you need to score text without any changes of scenery. With score 5 you need to score text with a moderate number of changes of the scenery. With score 9 you need to score text with several changes of the scenery. Answer only with a number. Here is the text in Italian: " |
| SOCIAL | "Evaluate how much an Italian text describes social interactions on a scale from 1 to 9. You should evaluate how much the situations include any kind of social interaction, either between two people or between more people, such as talking with somebody, yelling at somebody, taking part to an activity with other people. With score 1 you need to score text without any semantic relatedness to social interactions. With score 5 you need to score text with a moderate semantic relatedness to social interactions. With score 9 you need to score text with an extreme semantic relatedness to social interactions. Answer only with a number. Here is the text in Italian: " |
| SPACE | "Evaluate how much an Italian text describes the surrounding environment and the space where the events take place on a scale from 1 to 9. You should evaluate how much the text includes details regarding the environments and the spaces, such as descriptions of buildings, rooms or landscapes. With score 1 you need to score text without any descriptions of the surrounding environment. With score 5 you need to score text with moderate descriptions of the surrounding environment. With score 9 you need to score text with several descriptions of the surrounding environment. Answer only with a number. Here is the text in Italian: " |
| TACTILE | "Evaluate how much an Italian text describes tactile experiences on a scale from 1 to 9. You should evaluate how much the situations include the touching, details that can only be perceived through the touch with any body parts, or activities that include touching or brushing. With score 1 you need to score text without any semantic relatedness to tactile experiences. With score 5 you need to score text with a moderate semantic relatedness to tactile experiences. With score 9 you need to score text with extreme semantic relatedness to tactile experiences. Answer only with a number. Here is the text in Italian: " |
| TIME | "Evaluate how much an Italian text describes the temporal and sequential aspects of the events on a scale from 1 to 9. You should evaluate how much the text includes details regarding the chronological aspects, the duration and the time coordinates of the events. With score 1 you need to score text without any temporal and sequential aspects. With score 5 you need to score text with moderate temporal and sequential aspects. With score 9 you need to score text with several temporal and sequential aspects. Answer only with a number. Here is the text in Italian: " |
| VALENCE | "Evaluate the emotional valence of an Italian text on a scale from 1 to 9. The scoring is not related to the emotional strength or intensity. The scoring only concerns the subjective perception of positive, negative, or neutral tone. With score 1 you need to score text with extremely negative emotions, such as sadness. With score 5 you need to score text with neutral emotions, neither positive or negative. With score 9 you need to score text with extremely positive emotions such as happiness. Answer only with a number. Here is the text in Italian: " |
| VISUAL | "Evaluate how much an Italian text describes visual experiences on a scale from 1 to 9. You should evaluate how much the situations include vision, details that can only be perceived through the eyes and the vision, or activities that include vision. With score 1 you need to score text without any semantic relatedness to visual experiences. With score 5 you need to score text with a moderate semantic relatedness to visual experiences. With score 9 you need to score text with extreme semantic relatedness to visual experiences. Answer only with a number. Here is the text in Italian: " |

***Table S1. AIs' Prompting for the evaluation of semantic dimensions.*** *In the left column, the dimension labels. In the right column, the prompting used, including definitions, examples and meaning of the Likert Scale values.*

Table S2

| Dimension | Full Model adj $R^2$ | p-value Full model | report-type coefficient | Cohen's d | p-value non-parametric coefficient | q-value coefficient |
|---|---|---|---|---|---|---|
| **AGENTIVITY** | 0.183 | < 0.00001 | -1.58, CI: -1.70 -1.46 | -1.31 | < 0.00001 | **< 0.00001** |
| **AROUSAL** | 0.096 | 0.00002 | 0.21, CI: 0.10 0.31 | 0.19 | 0.01140 | **0.01216** |
| **AUDITORY** | 0.044 | < 0.00001 | 0.42, CI: 0.30 0.55 | 0.29 | < 0.00001 | **< 0.00001** |
| **BIZARRENESS** | 0.426 | < 0.00001 | 2.85, CI: 2.73 2.98 | 1.88 | < 0.00001 | **< 0.00001** |
| **BODY** | 0.076 | < 0.00001 | -0.73, CI: -0.85 -0.61 | -0.58 | < 0.00001 | **< 0.00001** |
| INCORPORATION | 0.014 | 0.11368 | -0.05, CI: -0.11 0.01 | -0.08 | 0.08945 | 0.08945 |
| **LIMITATIONS** | 0.106 | < 0.00001 | 0.82, CI: 0.69 0.95 | 0.54 | < 0.00001 | **< 0.00001** |
| **THOUGHT** | 0.346 | < 0.00001 | -1.77, CI: -1.88 -1.66 | -1.20 | < 0.00001 | **< 0.00001** |
| **MOVEMENTS** | 0.076 | < 0.00001 | 0.83, CI: 0.70 0.96 | 0.56 | < 0.00001 | **< 0.00001** |
| **SETTINGS** | 0.177 | < 0.00001 | 1.29, CI: 1.18 1.40 | 0.99 | < 0.00001 | **< 0.00001** |
| **SOCIAL** | 0.129 | < 0.00001 | 1.40, CI: 1.22 1.58 | 0.69 | < 0.00001 | **< 0.00001** |
| **SPACE** | 0.211 | < 0.00001 | 1.40, CI: 1.30 1.51 | 1.20 | < 0.00001 | **< 0.00001** |
| **TACTILE** | 0.064 | < 0.00001 | 0.49, CI: 0.40 0.58 | 0.60 | < 0.00001 | **< 0.00001** |
| **TIME** | 0.173 | < 0.00001 | -1.20, CI: -1.33 -1.08 | -0.72 | < 0.00001 | **< 0.00001** |
| **VALENCE** | 0.063 | < 0.00001 | -0.49, CI: -0.59 -0.38 | -0.43 | < 0.00001 | **< 0.00001** |
| **VISUAL** | 0.190 | < 0.00001 | 1.52, CI: 1.38 1.66 | 1.08 | < 0.00001 | **< 0.00001** |

***Table S2. Performance measures of the GLME model for the prediction of semantic dimensions across vigilance states in the discovery dataset.*** *The vigilance state (report-type coefficient) was used as a regressor of interest; age, sex, education level, and the BADA score were used as covariates of no interest; q<0.05, False discovery Rate -FDR- correction. Dimension labels in bold are those surviving FDR correction. Cohen's d refers to the effect of the report-type coefficient.*

Table S3

| | High-level Domain | Domain | Discovery dataset, dream reports | Discovery dataset, wakefulness reports | Test dataset, dream reports |
|---|---|---|---|---|---|
| | Community-centered | Toponyms | 19.03 ± 32.69 % | 16.05 ± 25.00 % | 19.52 ± 25.49 % |
| | Community-centered | Healthcare | 8.53 ± 14.05 % | 21.75 ± 25.48 % | 6.07 ± 14.42 % |
| | Community-centered | Jobs | 32.66 ± 27.40 % | 16.11 ± 25.00 % | 54.33 ± 33.47 % |
| | Community-centered | Drama | 60.53 ± 32.00 % | 65.39 ± 42.18 % | 85.38 ± 21.16 % |
| | Community-centered | Society | 20.42 ± 33.33 % | 13.89 ± 23.08 % | 14.78 ± 20.58 % |
| | Community-centered | Humanities | 11.30 ± 19.69 % | 28.64 ± 34.95 % | 14.29 ± 25.68 % |
| | Community-centered | Timing | 26.63 ± 24.72 % | 60.60 ± 40.95 % | 38.73 ± 31.47 % |
| | Community-centered | Technology | 7.93 ± 12.50 % | 17.67 ± 25.00 % | 11.54 ± 22.16 % |
| | Community-centered | Education | 22.29 ± 24.81 % | 31.76 ± 34.88 % | 34.66 ± 28.68 % |
| | Community-centered | Communication | 19.22 ± 29.64 % | 26.10 ± 28.46 % | 9.90 ± 15.80 % |
| | Community-centered | Slang | 5.56 ± 10.00 % | 8.06 ± 12.15 % | 7.44 ± 18.08 % |
| | Environment-centered | Food | 12.14 ± 18.18 % | 16.86 ± 28.25 % | 11.04 ± 18.78 % |
| | Environment-centered | Objects | 13.53 ± 22.22 % | 6.60 ± 11.11 % | 3.49 ± 9.89 % |
| | Environment-centered | Matter | 9.67 ± 16.67 % | 5.62 ± 9.09 % | 4.94 ± 12.13 % |
| | Environment-centered | Locations | 27.58 ± 28.89 % | 14.02 ± 22.22 % | 18.74 ± 24.41 % |
| | Environment-centered | Geometry | 24.25 ± 28.46 % | 10.06 ± 16.67 % | 16.16 ± 19.50 % |
| | Environment-centered | Nature | 29.80 ± 28.27 % | 21.13 ± 33.33 % | 47.86 ± 33.91 % |
| | Environment-centered | Measurements | 6.35 ± 11.11 % | 10.26 ± 16.67 % | 3.44 ± 9.90 % |
| | Environment-centered | Architecture | 20.16 ± 30.77 % | 7.34 ± 12.50 % | 6.02 ± 13.14 % |
| | Environment-centered | Animals | 18.44 ± 32.69 % | 8.74 ± 12.50 % | 20.63 ± 25.36 % |
| | Environment-centered | Appearance | 19.94 ± 30.00 % | 7.83 ± 12.15 % | 24.59 ± 27.83 % |
| | Environment-centered | Transportation | 19.14 ± 30.58 % | 12.04 ± 20.00 % | 13.27 ± 22.96 % |
| | Individual-centered | DOMAIN2 | 10.13 ± 16.67 % | 16.21 ± 25.00 % | 0.14 ± 1.24 % |
| | Individual-centered | DOMAIN3 | 15.49 ± 25.00 % | 13.65 ± 22.22 % | 18.27 ± 24.48 % |
| | Individual-centered | DOMAIN4 | 18.79 ± 30.77 % | 28.17 ± 25.71 % | 30.73 ± 28.83 % |
| | Individual-centered | Reactions | 14.16 ± 22.22 % | 17.34 ± 29.64 % | 32.71 ± 30.28 % |
| | Individual-centered | DOMAIN12 | 9.83 ± 16.35 % | 6.59 ± 11.11 % | 9.39 ± 21.73 % |
| | Individual-centered | Thriller | 21.89 ± 33.33 % | 9.68 ± 16.67 % | 10.61 ± 21.55 % |
| | Individual-centered | Fantasy | 7.50 ± 14.29 % | 10.18 ± 16.67 % | 15.02 ± 22.91 % |
| | Individual-centered | Concerns | 14.97 ± 22.22 % | 23.83 ± 33.33 % | 18.04 ± 24.04 % |
| | Individual-centered | Conflicts | 18.30 ± 30.00 % | 15.88 ± 25.00 % | 6.76 ± 12.89 % |
| | Individual-centered | Transaction | 16.61 ± 25.00 % | 14.76 ± 22.86 % | 5.81 ± 11.19 % |

***Table S3. Absolute frequencies of lexical domains.*** *Absolute frequencies of domain occurrences in dream and wakefulness reports, with standard deviations across individuals in the Discovery Dataset, and absolute frequencies in the Test Dataset ($q < 0.05$). In the first column, high-level semantic categories. In the second column, the lexical domain labels, as assigned by the raters.*

Table S4

| | Domain | Full Model adj $R^2$ | p-value Full model | report-type coefficient | Cohen's d | p-value non-parametric coefficient | q-value coefficient |
|---|---|---|---|---|---|---|---|
| | **DOMAIN2** | 0.066 | < 0.00001 | -0.53, CI: -0.75 -0.32 | -0.27 | 0.00005 | **0.00008** |
| | DOMAIN3 | 0.010 | 0.00339 | 0.11, CI: -0.08 0.30 | 0.09 | 0.30674 | 0.31663 |
| | **DOMAIN4** | 0.023 | < 0.00001 | -0.53, CI: -0.69 -0.37 | -0.37 | < 0.00001 | **< 0.00001** |
| | **Food** | 0.005 | 0.00047 | -0.37, CI: -0.56 -0.18 | -0.22 | 0.00129 | **0.00165** |
| | **Objects** | 0.047 | < 0.00001 | 0.80, CI: 0.57 1.02 | 0.40 | < 0.00001 | **< 0.00001** |
| | **Reactions** | 0.015 | 0.37539 | -0.21, CI: -0.39 -0.02 | -0.16 | 0.03753 | **0.04289** |
| | **DOMAIN12** | 0.015 | < 0.00001 | 0.50, CI: 0.24 0.75 | 0.19 | 0.00006 | **0.00008** |
| | **Matter** | 0.007 | 0.00007 | 0.56, CI: 0.31 0.82 | 0.31 | < 0.00001 | **< 0.00001** |
| | Toponyms | 0.034 | 0.00966 | 0.18, CI: 0.00 0.36 | 0.12 | 0.13689 | 0.15106 |
| | **Thriller** | 0.038 | < 0.00001 | 0.99, CI: 0.79 1.19 | 0.54 | < 0.00001 | **< 0.00001** |
| | **Healthcare** | 0.054 | < 0.00001 | -1.17, CI: -1.38 -0.97 | -0.64 | < 0.00001 | **< 0.00001** |
| | **Locations** | 0.061 | < 0.00001 | 0.84, CI: 0.67 1.02 | 0.53 | < 0.00001 | **< 0.00001** |
| | **Geometry** | 0.051 | < 0.00001 | 1.00, CI: 0.81 1.19 | 0.67 | < 0.00001 | **< 0.00001** |
| | **Nature** | 0.034 | < 0.00001 | 0.48, CI: 0.32 0.64 | 0.34 | < 0.00001 | **< 0.00001** |
| | **Fantasy** | 0.001 | 0.08989 | -0.32, CI: -0.55 -0.08 | -0.15 | 0.01126 | **0.01335** |
| | **Measurements** | 0.007 | 0.00003 | -0.56, CI: -0.82 -0.30 | -0.22 | < 0.00001 | **< 0.00001** |
| | **Architecture** | 0.032 | < 0.00001 | 1.08, CI: 0.86 1.29 | 0.61 | < 0.00001 | **< 0.00001** |
| | **Animals** | 0.071 | < 0.00001 | 0.93, CI: 0.72 1.14 | 0.50 | < 0.00001 | **< 0.00001** |
| | **Concerns** | 0.035 | < 0.00001 | -0.62, CI: -0.79 -0.44 | -0.34 | < 0.00001 | **< 0.00001** |
| | **Appearance** | 0.074 | < 0.00001 | 1.13, CI: 0.92 1.34 | 0.54 | < 0.00001 | **< 0.00001** |
| | **Jobs** | 0.046 | < 0.00001 | 0.97, CI: 0.80 1.14 | 0.60 | < 0.00001 | **< 0.00001** |
| | Conflicts | 0.013 | 0.47031 | 0.16, CI: -0.02 0.34 | 0.11 | 0.15134 | 0.16143 |
| | **Drama** | 0.072 | < 0.00001 | -0.25, CI: -0.40 -0.11 | -0.14 | 0.01096 | **0.01335** |
| | **Society** | 0.027 | < 0.00001 | 0.48, CI: 0.30 0.66 | 0.28 | < 0.00001 | **< 0.00001** |
| | **Humanities** | 0.098 | < 0.00001 | -1.20, CI: -1.39 -1.02 | -0.69 | < 0.00001 | **< 0.00001** |
| | **Transportation** | 0.010 | < 0.00001 | 0.51, CI: 0.32 0.70 | 0.33 | < 0.00001 | **< 0.00001** |
| | **Timing** | 0.167 | < 0.00001 | -1.54, CI: -1.69 -1.39 | -1.03 | < 0.00001 | **< 0.00001** |
| | **Technology** | 0.022 | < 0.00001 | -0.86, CI: -1.07 -0.65 | -0.48 | < 0.00001 | **< 0.00001** |
| | **Education** | 0.047 | < 0.00001 | -0.47, CI: -0.63 -0.32 | -0.33 | < 0.00001 | **< 0.00001** |
| | Transaction | 0.003 | 0.77129 | 0.07, CI: -0.11 0.25 | 0.09 | 0.45791 | 0.45791 |
| | **Communication** | 0.032 | < 0.00001 | -0.46, CI: -0.62 -0.30 | -0.25 | < 0.00001 | **< 0.00001** |
| | **Slang** | 0.015 | < 0.00001 | -0.46, CI: -0.73 -0.20 | -0.16 | 0.00108 | **0.00144** |

***Table S4. Performance measures of the GLME model for the prediction of lexical domains across vigilance states in the discovery dataset.*** *The vigilance state (report-type coefficient) was used as a regressor of interest; age, sex, education level, and the BADA score were used as covariates of no interest; q<0.05, False discovery Rate -FDR- correction. Domain labels in bold are those surviving FDR correction. Cohen's d refers to the effect of the report-type coefficient.*

Table S5

| Dimension | Full Model adj $R^2$ | p-value Full model | coefficient name | coefficient beta | p-value coefficient | q-value coefficient |
|---|---|---|---|---|---|---|
| AGENTIVITY | 0.183 | < 0.00001 | | | | |
| | | | report_type_1:Sex_1 | 0.33, CI: 0.05 0.61 | 0.02193 | 0.49897 |
| AROUSAL | 0.102 | < 0.00001 | | | | |
| | | | Sex_1 | -0.30, CI: -0.55 -0.06 | 0.01608 | 0.10853 |
| | | | Education | -0.05, CI: -0.09 -0.00 | 0.03327 | 0.17966 |
| | | | WC_BADA | 0.53, CI: 0.10 0.96 | 0.01557 | 0.10853 |
| | | | report_type_1:Education | 0.05, CI: 0.01 0.09 | 0.01544 | 0.10853 |
| | | | **report_type_1:ATD** | **0.05, CI: 0.02 0.08** | **0.00166** | **0.04492** |
| AUDITORY | 0.047 | < 0.00001 | | | | |
| | | | report_type_1:Age | -0.02, CI: -0.03 -0.00 | 0.00903 | 0.24384 |
| | | | report_type_1:WC_BADA | -0.72, CI: -1.36 -0.09 | 0.02610 | 0.35231 |
| BIZARRENESS | 0.433 | < 0.00001 | | | | |
| | | | report_type_1 | 2.85, CI: 0.56 5.14 | 0.01474 | 0.07961 |
| | | | report_type_1:Age | -0.01, CI: -0.03 -0.00 | 0.03043 | 0.13692 |
| | | | **report_type_1:PSQI** | **-0.09, CI: -0.15 -0.03** | **0.00277** | **0.02496** |
| | | | **report_type_1:ATD** | **0.07, CI: 0.03 0.10** | **0.00022** | **0.00296** |
| | | | **report_type_1:MW** | **0.27, CI: 0.16 0.38** | **< 0.00001** | **0.00005** |
| | | | **report_type_1:SCWT** | **-0.03, CI: -0.05 -0.01** | **0.00691** | **0.04665** |
| BODY | 0.076 | < 0.00001 | | | | |
| | | | report_type_1 | -3.17, CI: -5.45 -0.90 | 0.00629 | 0.08496 |
| | | | Age | -0.02, CI: -0.03 -0.01 | 0.00200 | 0.05396 |
| | | | STAI | -0.02, CI: -0.03 -0.00 | 0.02561 | 0.23052 |
| LIMITATIONS | 0.109 | < 0.00001 | | | | |
| | | | Sex_1 | -0.41, CI: -0.70 -0.12 | 0.00503 | 0.08745 |
| | | | ATD | -0.04, CI: -0.07 -0.00 | 0.03834 | 0.25881 |
| | | | WC_BADA | 0.68, CI: 0.17 1.20 | 0.00972 | 0.08745 |
| | | | report_type_1:ATD | 0.05, CI: 0.01 0.09 | 0.00660 | 0.08745 |
| THOUGHT | 0.352 | < 0.00001 | | | | |
| | | | Sex_1 | -0.33, CI: -0.64 -0.01 | 0.04030 | 0.13952 |
| | | | Age | -0.02, CI: -0.03 -0.01 | 0.00558 | 0.07526 |
| | | | STAI | 0.02, CI: 0.00 0.04 | 0.04134 | 0.13952 |
| | | | MW | 0.15, CI: 0.03 0.27 | 0.01820 | 0.13952 |
| | | | MEQ | -0.02, CI: -0.03 -0.00 | 0.03820 | 0.13952 |
| | | | **WC_BADA** | **0.93, CI: 0.46 1.40** | **0.00012** | **0.00314** |
| | | | report_type_1:STAI | -0.02, CI: -0.03 -0.00 | 0.04995 | 0.14984 |
| | | | report_type_1:BSRT | -0.04, CI: -0.08 -0.01 | 0.02182 | 0.13952 |
| | | | report_type_1:MEQ | 0.01, CI: 0.00 0.02 | 0.03483 | 0.13952 |
| MOVEMENTS | 0.079 | < 0.00001 | | | | |
| | | | MEQ | 0.02, CI: 0.00 0.03 | 0.00898 | 0.08082 |
| | | | report_type_1:ATD | 0.05, CI: 0.02 0.09 | 0.00386 | 0.05217 |

| | | | | | | |
|---|---|---|---|---|---|---|
| | | | report_type_1:MW | 0.12, CI: 0.00 0.24 | 0.04276 | 0.28864 |
| | | | report_type_1:MEQ | -0.02, CI: -0.03 -0.01 | 0.00385 | 0.05217 |
| SETTINGS | 0.185 | < 0.00001 | | | | |
| | | | report_type_1:ATD | 0.04, CI: 0.01 0.08 | 0.00482 | 0.06501 |
| | | | **report_type_1:MW** | **0.23, CI: 0.14 0.33** | **< 0.00001** | **0.00008** |
| | | | report_type_1:VVIQ | 0.01, CI: 0.00 0.02 | 0.04540 | 0.40857 |
| SOCIAL | 0.135 | < 0.00001 | | | | |
| | | | Sex_1 | -0.59, CI: -0.98 -0.20 | 0.00336 | 0.09082 |
| | | | SCWT | -0.04, CI: -0.07 -0.01 | 0.00729 | 0.09846 |
| | | | WC_BADA | 0.83, CI: 0.12 1.53 | 0.02145 | 0.09880 |
| | | | report_type_1:Sex_1 | 0.48, CI: 0.07 0.89 | 0.02172 | 0.09880 |
| | | | report_type_1:Age | -0.02, CI: -0.04 -0.00 | 0.02016 | 0.09880 |
| | | | report_type_1:MW | 0.17, CI: 0.01 0.33 | 0.04170 | 0.14630 |
| | | | report_type_1:BSRT | 0.06, CI: 0.00 0.11 | 0.04335 | 0.14630 |
| | | | report_type_1:WC_BADA | -1.07, CI: -1.99 -0.16 | 0.02196 | 0.09880 |
| SPACE | 0.217 | < 0.00001 | | | | |
| | | | **report_type_1:ATD** | **0.07, CI: 0.04 0.10** | **< 0.00001** | **0.00017** |
| | | | report_type_1:MW | 0.11, CI: 0.02 0.21 | 0.02284 | 0.26923 |
| | | | report_type_1:BSRT | 0.04, CI: 0.00 0.07 | 0.02991 | 0.26923 |
| TACTILE | 0.064 | < 0.00001 | | | | |
| | | | MEQ | 0.01, CI: 0.00 0.02 | 0.00256 | 0.06923 |
| | | | report_type_1:WC_BADA | 0.49, CI: 0.03 0.95 | 0.03672 | 0.49573 |
| TIME | 0.178 | < 0.00001 | | | | |
| | | | Age | -0.02, CI: -0.03 -0.01 | 0.00297 | 0.08019 |
| | | | VVIQ | 0.01, CI: 0.00 0.03 | 0.01842 | 0.12436 |
| | | | report_type_1:ATD | 0.05, CI: 0.01 0.08 | 0.01134 | 0.10209 |
| | | | report_type_1:MW | 0.15, CI: 0.03 0.26 | 0.01042 | 0.10209 |
| VALENCE | 0.068 | < 0.00001 | | | | |
| | | | ATD | 0.03, CI: 0.00 0.05 | 0.03019 | 0.19031 |
| | | | BSRT | -0.03, CI: -0.06 -0.00 | 0.02966 | 0.19031 |
| | | | VVIQ | 0.01, CI: 0.00 0.02 | 0.04276 | 0.19031 |
| | | | report_type_1:PSQI | 0.07, CI: 0.02 0.12 | 0.00727 | 0.19031 |
| | | | report_type_1:BSRT | 0.04, CI: 0.01 0.07 | 0.01463 | 0.19031 |
| | | | report_type_1:SCWT | 0.02, CI: 0.00 0.03 | 0.03620 | 0.19031 |
| VISUAL | 0.193 | < 0.00001 | | | | |
| | | | Sex_1 | -0.36, CI: -0.66 -0.06 | 0.01991 | 0.17921 |
| | | | report_type_1:Sex_1 | 0.43, CI: 0.12 0.74 | 0.00712 | 0.09612 |
| | | | **report_type_1:ATD** | **0.07, CI: 0.03 0.10** | **0.00079** | **0.02136** |

***Table S5. Performance measures of the GLME model for the prediction of semantic dimensions based on individual psychological variables in the discovery dataset.*** *The GLME model included psychological variables and their interaction with vigilance states (report_type) as regressors of interest and age, sex, education level, and the BADA score as covariates; q<0.05, False discovery Rate -FDR- correction. Significant effects after FDR correction are reported in bold.*

Table S6

| | Domain | Full Model adj $R^2$ | p-value Full model | coefficient name | coefficient beta | p-value coefficient | q-value coefficient |
|---|---|---|---|---|---|---|---|
| | DOMAIN2 | 0.067 | 0.00005 | | | | |
| | | | | Sex_1 | 0.48, CI: 0.07 0.89 | 0.02106 | 0.18955 |
| | | | | Age | 0.02, CI: 0.00 0.04 | 0.01312 | 0.17707 |
| | | | | report_type_1:Sex_1 | -0.56, CI: -1.07 -0.06 | 0.02899 | 0.19570 |
| | | | | report_type_1:VVIQ | 0.03, CI: 0.01 0.05 | 0.00638 | 0.17214 |
| | DOMAIN4 | 0.023 | 0.00001 | | | | |
| | Food | 0.020 | 0.04967 | | | | |
| | | | | STAI | -0.02, CI: -0.04 -0.00 | 0.02337 | 0.31552 |
| | | | | report_type_1:STAI | 0.03, CI: 0.00 0.06 | 0.02041 | 0.31552 |
| | Objects | 0.027 | < 0.00001 | | | | |
| | | | | MW | 0.20, CI: 0.03 0.36 | 0.02102 | 0.18917 |
| | | | | **ROCFr** | **-0.07, CI: -0.11 -0.03** | **0.00034** | **0.00564** |
| | | | | **report_type_1:ROCFr** | **0.09, CI: 0.04 0.14** | **0.00042** | **0.00564** |
| | Reactions | 0.016 | 0.34785 | | | | |
| | | | | report_type_1:Sex_1 | -0.44, CI: -0.88 -0.01 | 0.04400 | 0.59405 |
| | | | | report_type_1:MW | -0.19, CI: -0.36 -0.02 | 0.02596 | 0.59405 |
| | DOMAIN12 | 0.016 | 0.00026 | | | | |
| | | | | **Age** | **-0.04, CI: -0.06 -0.02** | **0.00064** | **0.01725** |
| | | | | ROCFr | 0.05, CI: 0.00 0.09 | 0.03856 | 0.50457 |
| | Matter | 0.014 | 0.00033 | | | | |
| | | | | **PSQI** | **-0.18, CI: -0.30 -0.06** | **0.00245** | **0.03306** |
| | | | | BSRT | 0.07, CI: 0.01 0.13 | 0.03294 | 0.25087 |
| | | | | WC_BADA | 1.02, CI: 0.06 1.97 | 0.03717 | 0.25087 |
| | | | | **report_type_1:PSQI** | **0.22, CI: 0.08 0.36** | **0.00193** | **0.03306** |
| | Thriller | 0.036 | < 0.00001 | | | | |
| | | | | Sex_1 | -0.42, CI: -0.80 -0.04 | 0.03022 | 0.29247 |
| | | | | Age | -0.02, CI: -0.03 -0.00 | 0.04070 | 0.29247 |
| | | | | BSRT | 0.06, CI: 0.01 0.11 | 0.02726 | 0.29247 |
| | | | | report_type_1:Sex_1 | 0.48, CI: 0.01 0.94 | 0.04333 | 0.29247 |
| | Healthcare | 0.042 | < 0.00001 | | | | |
| | | | | report_type_1:Age | 0.02, CI: 0.00 0.04 | 0.03491 | 0.47133 |
| | | | | report_type_1:MEQ | -0.02, CI: -0.05 -0.00 | 0.02742 | 0.47133 |
| | Locations | 0.067 | < 0.00001 | | | | |
| | | | | MW | -0.16, CI: -0.30 -0.01 | 0.03198 | 0.28783 |
| | | | | BSRT | 0.07, CI: 0.02 0.12 | 0.00499 | 0.09158 |
| | | | | report_type_1:ATD | 0.05, CI: 0.00 0.10 | 0.04628 | 0.31236 |
| | | | | report_type_1:MW | 0.22, CI: 0.06 0.38 | 0.00678 | 0.09158 |
| | Geometry | 0.056 | < 0.00001 | | | | |
| | | | | **report_type_1:ATD** | **0.09, CI: 0.04 0.15** | **0.00060** | **0.01627** |
| | Nature | 0.036 | 0.00005 | | | | |

| | | | | | | | |
|---|---|---|---|---|---|---|---|
| | | | | MEQ | 0.02, CI: 0.00 0.03 | 0.01670 | 0.22550 |
| | | | | **report_type_1:ATD** | **0.07, CI: 0.03 0.12** | **0.00176** | **0.04748** |
| | | | | report_type_1:MEQ | -0.02, CI: -0.03 -0.00 | 0.04567 | 0.41107 |
| | Fantasy | 0.005 | 0.04715 | | | | |
| | | | | BSRT | 0.06, CI: 0.01 0.10 | 0.02616 | 0.35309 |
| | | | | report_type_1:Age | -0.03, CI: -0.05 -0.00 | 0.02398 | 0.35309 |
| | Measurements | 0.006 | 0.01810 | | | | |
| | Architecture | 0.051 | < 0.00001 | | | | |
| | Animals | 0.080 | < 0.00001 | | | | |
| | | | | BSRT | 0.08, CI: 0.02 0.14 | 0.01202 | 0.32445 |
| | Concerns | 0.035 | < 0.00001 | | | | |
| | | | | Sex_1 | 0.37, CI: 0.08 0.67 | 0.01361 | 0.18374 |
| | | | | STAI | 0.02, CI: 0.00 0.04 | 0.03784 | 0.34054 |
| | | | | WC_BADA | 0.76, CI: 0.17 1.35 | 0.01129 | 0.18374 |
| | Appearance | 0.082 | < 0.00001 | | | | |
| | | | | **report_type_1** | **-5.82, CI: -9.87 -1.76** | **0.00494** | **0.04854** |
| | | | | Sex_1 | -0.49, CI: -0.98 -0.01 | 0.04661 | 0.17362 |
| | | | | Age | -0.02, CI: -0.05 -0.00 | 0.01971 | 0.11271 |
| | | | | **PSQI** | **-0.15, CI: -0.26 -0.05** | **0.00539** | **0.04854** |
| | | | | MW | 0.21, CI: 0.03 0.39 | 0.02455 | 0.11271 |
| | | | | report_type_1:Education | 0.10, CI: 0.01 0.18 | 0.02505 | 0.11271 |
| | | | | **report_type_1:PSQI** | **0.17, CI: 0.06 0.29** | **0.00331** | **0.04854** |
| | Jobs | 0.050 | < 0.00001 | | | | |
| | | | | Age | 0.02, CI: 0.00 0.03 | 0.00723 | 0.06508 |
| | | | | report_type_1:Sex_1 | 0.56, CI: 0.16 0.95 | 0.00543 | 0.06508 |
| | | | | **report_type_1:Age** | **-0.03, CI: -0.04 -0.01** | **0.00110** | **0.02973** |
| | | | | report_type_1:MEQ | -0.02, CI: -0.04 -0.00 | 0.04027 | 0.27180 |
| | Drama | 0.079 | < 0.00001 | | | | |
| | | | | Sex_1 | -0.42, CI: -0.72 -0.12 | 0.00653 | 0.12416 |
| | | | | Age | -0.01, CI: -0.03 -0.00 | 0.04102 | 0.22015 |
| | | | | STAI | 0.02, CI: 0.00 0.04 | 0.02903 | 0.22015 |
| | | | | PSQI | -0.08, CI: -0.15 -0.02 | 0.00920 | 0.12416 |
| | | | | WC_BADA | 0.60, CI: 0.02 1.18 | 0.04103 | 0.22015 |
| | | | | report_type_1:MEQ | 0.02, CI: 0.00 0.03 | 0.04892 | 0.22015 |
| | Society | 0.031 | 0.00010 | | | | |
| | | | | STAI | -0.03, CI: -0.05 -0.01 | 0.01050 | 0.09450 |
| | | | | WC_BADA | 1.05, CI: 0.35 1.75 | 0.00318 | 0.08576 |
| | | | | report_type_1:WC_BADA | -1.22, CI: -2.12 -0.32 | 0.00796 | 0.09450 |
| | Humanities | 0.112 | < 0.00001 | | | | |
| | | | | Sex_1 | 0.42, CI: 0.10 0.73 | 0.01016 | 0.09144 |
| | | | | Age | -0.01, CI: -0.03 -0.00 | 0.04734 | 0.27520 |
| | | | | report_type_1:Sex_1 | -0.66, CI: -1.10 -0.22 | 0.00312 | 0.05163 |
| | | | | report_type_1:PSQI | 0.13, CI: 0.04 0.22 | 0.00382 | 0.05163 |
| | Transportation | 0.018 | 0.00216 | | | | |

| | | | | | | |
|---|---|---|---|---|---|---|
| Timing | 0.179 | < 0.00001 | | | | |
| | | | report_type_1 | -3.74, CI: -6.49 -1.00 | 0.00751 | 0.10133 |
| | | | MW | -0.11, CI: -0.22 -0.00 | 0.04425 | 0.29869 |
| | | | report_type_1:Education | 0.09, CI: 0.03 0.15 | 0.00192 | 0.05195 |
| | | | report_type_1:ATD | 0.05, CI: 0.01 0.09 | 0.01805 | 0.16241 |
| Technology | 0.023 | < 0.00001 | | | | |
| | | | Education | 0.06, CI: 0.01 0.11 | 0.01440 | 0.19435 |
| | | | SCWT | -0.02, CI: -0.04 -0.00 | 0.04057 | 0.29351 |
| | | | WC_BADA | 0.61, CI: 0.02 1.21 | 0.04406 | 0.29351 |
| | | | report_type_1:WC_BADA | -1.29, CI: -2.31 -0.26 | 0.01427 | 0.19435 |
| Education | 0.046 | < 0.00001 | | | | |
| | | | **Age** | **-0.02, CI: -0.04 -0.01** | **0.00022** | **0.00593** |
| | | | report_type_1:VVIQ | -0.02, CI: -0.03 -0.00 | 0.03059 | 0.41295 |
| Communication | 0.035 | < 0.00001 | | | | |
| | | | **Age** | **-0.03, CI: -0.04 -0.01** | **0.00013** | **0.00352** |
| | | | MEQ | -0.02, CI: -0.03 -0.00 | 0.01911 | 0.17203 |
| | | | report_type_1:Education | 0.07, CI: 0.00 0.13 | 0.04110 | 0.27743 |
| | | | **report_type_1:MEQ** | **0.03, CI: 0.01 0.04** | **0.00353** | **0.04769** |
| Slang | 0.017 | < 0.00001 | | | | |
| | | | **Sex_1** | **0.64, CI: 0.27 1.02** | **0.00085** | **0.01454** |
| | | | **Age** | **-0.03, CI: -0.06 -0.01** | **0.00145** | **0.01454** |
| | | | PSQI | -0.12, CI: -0.21 -0.03 | 0.01076 | 0.07261 |
| | | | ATD | -0.05, CI: -0.10 -0.01 | 0.02683 | 0.14486 |
| | | | report_type_1:Sex_1 | -0.64, CI: -1.24 -0.04 | 0.03706 | 0.16678 |
| | | | **report_type_1:MW** | **-0.39, CI: -0.63 -0.15** | **0.00162** | **0.01454** |

***Table S6. Performance measures of the GLME model for the prediction of lexical domains based on individual psychological variables in the discovery dataset.*** *The GLME model included psychological variables and their interaction with vigilance states (report_type) as regressors of interest and age, sex, education level, and the BADA score as covariates; q<0.05, False discovery Rate -FDR- correction. Significant effects after FDR correction are reported in bold.*

Table S7

| Dimension | Full Model adj $R^2$ | p-value Full model | coefficient name | coefficient beta | p-value coefficient | q-value coefficient |
|---|---|---|---|---|---|---|
| AGENTIVITY | 0.013 | 0.11558 | | | | |
| AROUSAL | 0.110 | 0.23799 | | | | |
| | | | PC4_report | 0.07, CI: 0.00 0.13 | 0.03539 | 0.60155 |
| AUDITORY | 0.050 | 0.28287 | | | | |
| | | | Age | -0.01, CI: -0.02 -0.00 | 0.02824 | 0.48008 |
| BIZARRENESS | 0.169 | 0.00001 | | | | |
| | | | Age | -0.02, CI: -0.04 -0.00 | 0.03236 | 0.13752 |
| | | | PSQI | -0.14, CI: -0.23 -0.05 | 0.00194 | 0.01652 |
| | | | MW | 0.35, CI: 0.18 0.51 | 0.00006 | 0.00094 |
| | | | SCWT | -0.03, CI: -0.06 -0.00 | 0.03218 | 0.13752 |
| BODY | 0.034 | 0.66715 | | | | |
| LIMITATIONS | 0.099 | 0.39999 | | | | |
| THOUGHT | 0.177 | 0.13254 | | | | |
| | | | WC_BADA | 0.92, CI: 0.26 1.59 | 0.00638 | 0.10854 |
| MOVEMENTS | 0.090 | 0.02157 | | | | |
| | | | ATD | 0.04, CI: 0.00 0.08 | 0.03887 | 0.33043 |
| | | | PC3_report | 0.11, CI: 0.02 0.19 | 0.01098 | 0.18659 |
| SETTINGS | 0.114 | 0.00010 | | | | |
| | | | MW | 0.22, CI: 0.09 0.35 | 0.00112 | 0.01899 |
| | | | **PC2_report** | **0.09, CI: 0.03 0.16** | **0.00274** | **0.02325** |
| SOCIAL | 0.076 | 0.00083 | | | | |
| | | | Age | -0.03, CI: -0.05 -0.01 | 0.00026 | 0.00438 |
| | | | PC1_report | 0.06, CI: 0.01 0.10 | 0.01041 | 0.08844 |
| SPACE | 0.139 | 0.00046 | | | | |
| | | | ATD | 0.07, CI: 0.03 0.11 | 0.00029 | 0.00496 |
| | | | BSRT | 0.05, CI: 0.01 0.10 | 0.01950 | 0.11051 |
| | | | PC2_report | 0.08, CI: 0.02 0.13 | 0.01182 | 0.10046 |
| TACTILE | 0.051 | 0.43745 | | | | |
| TIME | 0.096 | 0.04319 | | | | |
| | | | PC2_report | 0.06, CI: 0.01 0.12 | 0.03168 | 0.53855 |
| VALENCE | 0.031 | 0.49050 | | | | |
| | | | PC4_report | -0.05, CI: -0.10 -0.00 | 0.03326 | 0.56540 |
| VISUAL | 0.122 | 0.00784 | | | | |
| | | | ATD | 0.07, CI: 0.03 0.12 | 0.00222 | 0.03767 |

***Table S7. Performance measures of the GLME model for the prediction of semantic dimensions in dream reports from the discovery dataset, based on sleep patterns.*** *The GLME model included all individual predictors that were found to be significant (uncorrected p-value < 0.05) in the previous analysis. For assessing the impact of sleep macrostructure, scores for the four actigraphic PCs were included as regressors of interest; q<0.05, False discovery Rate -FDR- correction. Significant effects for actigraphic PCs after FDR correction are reported in bold.*

Table S8

| | Domain | Full Model adj $R^2$ | p-value Full model | coefficient name | coefficient beta | p-value coefficient | q-value coefficient |
|---|---|---|---|---|---|---|---|
| | DOMAIN2 | 0.059 | 0.37824 | | | | |
| | Food | 0.010 | 0.37056 | | | | |
| | Objects | 0.006 | 0.01581 | | | | |
| | | | | MW | 0.20, CI: 0.08 0.33 | 0.00100 | 0.01699 |
| | Reactions | -0.000 | 0.45481 | | | | |
| | DOMAIN12 | 0.010 | 0.01151 | | | | |
| | | | | Age | -0.03, CI: -0.05 -0.01 | 0.00061 | 0.01044 |
| | Matter | -0.002 | 0.79193 | | | | |
| | Thriller | 0.026 | 0.34324 | | | | |
| | Healthcare | 0.007 | 0.00459 | | | | |
| | | | | Age | 0.02, CI: 0.00 0.03 | 0.04305 | 0.24398 |
| | | | | MEQ | -0.04, CI: -0.06 -0.02 | 0.00024 | 0.00406 |
| | | | | PC4_report | -0.13, CI: -0.26 -0.01 | 0.03594 | 0.24398 |
| | Locations | 0.038 | 0.01283 | | | | |
| | | | | ATD | 0.04, CI: 0.00 0.08 | 0.03888 | 0.16524 |
| | | | | BSRT | 0.04, CI: 0.00 0.08 | 0.03280 | 0.16524 |
| | | | | PC2_report | 0.08, CI: 0.02 0.15 | 0.01265 | 0.16524 |
| | | | | PC4_report | -0.09, CI: -0.17 -0.01 | 0.03643 | 0.16524 |
| | Geometry | 0.034 | 0.02059 | | | | |
| | | | | ATD | 0.05, CI: 0.02 0.09 | 0.00519 | 0.08826 |
| | | | | PC3_report | 0.09, CI: 0.01 0.17 | 0.02227 | 0.18925 |
| | Nature | 0.040 | 0.11205 | | | | |
| | | | | ATD | 0.05, CI: 0.01 0.08 | 0.01303 | 0.22150 |
| | Fantasy | -0.001 | 0.56986 | | | | |
| | Animals | 0.065 | 0.80568 | | | | |
| | Concerns | 0.001 | 0.30704 | | | | |
| | | | | PC3_report | -0.10, CI: -0.19 -0.01 | 0.02545 | 0.43268 |
| | Appearance | 0.075 | 0.07281 | | | | |
| | | | | Sex | -0.37, CI: -0.74 -0.00 | 0.04790 | 0.41799 |
| | | | | PC2_report | 0.08, CI: 0.00 0.15 | 0.04918 | 0.41799 |
| | Jobs | 0.016 | 0.00160 | | | | |
| | | | | Age | -0.01, CI: -0.02 -0.00 | 0.00862 | 0.07325 |
| | | | | PC2_report | -0.08, CI: -0.14 -0.02 | 0.00763 | 0.07325 |
| | Drama | 0.048 | 0.00013 | | | | |
| | | | | Sex | -0.64, CI: -0.91 -0.36 | < 0.00001 | 0.00010 |
| | | | | Age | -0.02, CI: -0.03 -0.01 | 0.00040 | 0.00336 |
| | Society | 0.025 | 0.29664 | | | | |
| | Humanities | 0.016 | 0.00015 | | | | |
| | | | | Age | -0.03, CI: -0.04 -0.01 | 0.00105 | 0.01783 |
| | | | | PSQI | 0.08, CI: 0.01 0.14 | 0.02103 | 0.17873 |

| | | | | | | |
|---|---|---|---|---|---|---|
| Timing | 0.009 | 0.00717 | | | | |
| | | | Education | 0.05, CI: 0.01 0.09 | 0.02127 | 0.18076 |
| | | | PC2_report | 0.06, CI: 0.00 0.12 | 0.04757 | 0.26959 |
| | | | PC3_report | 0.10, CI: 0.03 0.17 | 0.00678 | 0.11530 |
| Technology | 0.001 | 0.27806 | | | | |
| Education | 0.012 | 0.13195 | | | | |
| Communication | 0.003 | 0.12507 | | | | |
| | | | MEQ | 0.01, CI: 0.00 0.03 | 0.03817 | 0.43460 |
| Slang | 0.006 | 0.02751 | | | | |
| | | | MW | -0.29, CI: -0.48 -0.10 | 0.00305 | 0.05178 |

***Table S8. Performance measures of the GLME model for the prediction of lexical domains in dream reports from the discovery dataset, based on sleep patterns.*** *The GLME model included all individual predictors that were found to be significant (uncorrected p-value < 0.05) in the previous analysis. For assessing the impact of sleep macrostructure, scores for the four actigraphic PCs were included as regressors of interest; q<0.05, False discovery Rate -FDR- correction. Significant effects for actigraphic PCs after FDR correction are reported in bold.*

Table S9

| Dimension | Full Model adj $R^2$ | p-value Full model | Dataset coefficient | Cohen's d | p-value coefficient | q-value coefficient |
|---|---|---|---|---|---|---|
| AGENTIVITY | 0.030 | < 0.00001 | 0.27, CI: 0.00 0.54 | 0.27 | 0.04657 | 0.09313 |
| **AROUSAL** | **0.164** | **< 0.00001** | **0.40, CI: 0.17 0.63** | **0.50** | **0.00060** | **0.00482** |
| AUDITORY | 0.057 | < 0.00001 | -0.17, CI: -0.45 0.12 | -0.19 | 0.25368 | 0.36881 |
| BIZARRENESS | 0.203 | < 0.00001 | 0.07, CI: -0.26 0.39 | -0.02 | 0.68309 | 0.78067 |
| **BODY** | **0.054** | **< 0.00001** | **0.34, CI: 0.09 0.60** | **0.36** | **0.00869** | **0.02800** |
| INCORPORATION | -0.001 | 0.63139 | -0.07, CI: -0.18 0.04 | -0.12 | 0.23750 | 0.36881 |
| **LIMITATIONS** | **0.143** | **< 0.00001** | **0.43, CI: 0.15 0.71** | **0.46** | **0.00281** | **0.01496** |
| THOUGHT | 0.214 | < 0.00001 | -0.16, CI: -0.41 0.09 | -0.03 | 0.19926 | 0.35423 |
| **MOVEMENTS** | **0.110** | **< 0.00001** | **0.41, CI: 0.07 0.74** | **0.35** | **0.01689** | **0.04504** |
| **SETTINGS** | **0.181** | **< 0.00001** | **0.50, CI: 0.22 0.78** | **0.41** | **0.00053** | **0.00482** |
| **SOCIAL** | **0.126** | **< 0.00001** | **0.47, CI: 0.12 0.82** | **0.48** | **0.00875** | **0.02800** |
| SPACE | 0.175 | < 0.00001 | 0.02, CI: -0.26 0.29 | -0.16 | 0.91321 | 0.91321 |
| TACTILE | 0.064 | < 0.00001 | -0.01, CI: -0.23 0.20 | -0.10 | 0.90531 | 0.91321 |
| TIME | 0.172 | < 0.00001 | 0.26, CI: 0.03 0.49 | 0.30 | 0.02966 | 0.06778 |
| VALENCE | 0.029 | 0.06388 | -0.06, CI: -0.27 0.15 | -0.20 | 0.58434 | 0.71919 |
| VISUAL | 0.150 | < 0.00001 | 0.17, CI: -0.13 0.46 | 0.09 | 0.27661 | 0.36881 |

***Table S9. Performance measures of the GLME model for the prediction of semantic dimensions across discovery and test datasets.*** *The GLME model included sex, age, educational level, and the average word count per participant as covariates of no interest; q<0.05, False discovery Rate -FDR- correction. Significant effects after FDR correction are reported in bold. Cohen's d refers to the effect of the dataset (Test – Discovery datasets).*

Table S10

| | Domain | Full Model adj $R^2$ | p-value Full model | Dataset coefficient | Cohen's d | p-value coefficient | q-value coefficient |
|---|---|---|---|---|---|---|---|
| | **DOMAIN2** | **0.120** | **0.00124** | **-3.29, CI: -5.30 -1.28** | **-0.79** | **0.00132** | **0.00325** |
| | DOMAIN3 | 0.009 | 0.07131 | -0.13, CI: -0.50 0.24 | 0.15 | 0.47861 | 0.54972 |
| | DOMAIN4 | 0.035 | < 0.00001 | 0.11, CI: -0.19 0.41 | 0.57 | 0.48100 | 0.54972 |
| | Food | 0.019 | < 0.00001 | -0.32, CI: -0.71 0.07 | -0.07 | 0.11223 | 0.14963 |
| | **Objects** | **0.056** | **< 0.00001** | **-1.86, CI: -2.44 -1.28** | **-0.70** | **< 0.00001** | **< 0.00001** |
| | **Reactions** | **0.023** | **< 0.00001** | **0.74, CI: 0.44 1.04** | **0.93** | **< 0.00001** | **< 0.00001** |
| | DOMAIN12 | 0.012 | 0.00110 | -0.47, CI: -0.92 -0.01 | -0.02 | 0.04558 | 0.07293 |
| | **Matter** | **0.023** | **< 0.00001** | **-0.78, CI: -1.29 -0.26** | **-0.36** | **0.00326** | **0.00651** |
| | Toponyms | 0.069 | < 0.00001 | -0.32, CI: -0.70 0.05 | 0.03 | 0.09397 | 0.13074 |
| | **Thriller** | **0.031** | **< 0.00001** | **-1.22, CI: -1.62 -0.83** | **-0.56** | **< 0.00001** | **< 0.00001** |
| | Healthcare | 0.003 | 0.09021 | -0.50, CI: -1.05 0.04 | -0.19 | 0.07140 | 0.10879 |
| | **Locations** | **0.065** | **< 0.00001** | **-0.51, CI: -0.84 -0.18** | **-0.39** | **0.00227** | **0.00484** |
| | **Geometry** | **0.072** | **< 0.00001** | **-0.78, CI: -1.11 -0.45** | **-0.43** | **< 0.00001** | **0.00002** |
| | **Nature** | **0.110** | **< 0.00001** | **0.45, CI: 0.18 0.72** | **0.69** | **0.00115** | **0.00307** |
| | **Fantasy** | **0.014** | **< 0.00001** | **0.56, CI: 0.16 0.95** | **0.50** | **0.00553** | **0.00984** |
| | Measurements | 0.002 | 0.12883 | -0.38, CI: -1.00 0.24 | -0.29 | 0.22883 | 0.28164 |
| | **Architecture** | **0.055** | **< 0.00001** | **-1.41, CI: -1.86 -0.96** | **-0.83** | **< 0.00001** | **< 0.00001** |
| | Animals | 0.083 | < 0.00001 | 0.09, CI: -0.30 0.47 | 0.12 | 0.65985 | 0.72811 |
| | **Concerns** | **0.005** | **0.00718** | **0.42, CI: 0.07 0.77** | **0.18** | **0.01783** | **0.03003** |
| | Appearance | 0.100 | < 0.00001 | -0.03, CI: -0.39 0.34 | 0.20 | 0.89127 | 0.92002 |
| | **Jobs** | **0.048** | **< 0.00001** | **0.86, CI: 0.58 1.15** | **0.88** | **< 0.00001** | **< 0.00001** |
| | **Conflicts** | **0.053** | **< 0.00001** | **-1.21, CI: -1.68 -0.75** | **-0.70** | **< 0.00001** | **< 0.00001** |
| | **Drama** | **0.118** | **< 0.00001** | **0.68, CI: 0.35 1.01** | **1.09** | **0.00006** | **0.00020** |
| | Society | 0.034 | < 0.00001 | -0.31, CI: -0.67 0.05 | -0.29 | 0.09249 | 0.13074 |
| | Humanities | 0.010 | 0.00117 | -0.32, CI: -0.73 0.09 | 0.18 | 0.12528 | 0.16036 |
| | **Transportation** | **0.028** | **< 0.00001** | **-0.58, CI: -0.95 -0.21** | **-0.30** | **0.00197** | **0.00449** |
| | **Timing** | **0.025** | **< 0.00001** | **0.53, CI: 0.25 0.82** | **0.51** | **0.00024** | **0.00071** |
| | Technology | 0.004 | 0.03167 | -0.07, CI: -0.53 0.40 | 0.27 | 0.77246 | 0.82396 |
| | **Education** | **0.021** | **0.00003** | **0.46, CI: 0.15 0.77** | **0.58** | **0.00360** | **0.00677** |
| | **Transaction** | **0.028** | **< 0.00001** | **-1.19, CI: -1.64 -0.74** | **-0.77** | **< 0.00001** | **< 0.00001** |
| | **Communication** | **0.009** | **0.00131** | **-0.75, CI: -1.13 -0.38** | **-0.55** | **0.00008** | **0.00027** |
| | Slang | 0.011 | 0.00061 | -0.02, CI: -0.52 0.47 | 0.14 | 0.92780 | 0.92780 |

***Table S10. Performance measures of the GLME model for the prediction of lexical domains across discovery and test datasets.*** *The GLME model included sex, age, educational level, and the average word count per participant as covariates of no interest; q<0.05, False discovery Rate -FDR- correction. Significant effects after FDR correction are reported in bold. Cohen's d refers to the effect of the dataset (Test – Discovery datasets).*

Table S11

| Dimension | Full Model adj $R^2$ | p-value Full model | coefficient name | coefficient beta | p-value coefficient | q-value coefficient |
|---|---|---|---|---|---|---|
| AGENTIVITY | 0.183 | < 0.00001 | | | | |
| AROUSAL | 0.098 | < 0.00001 | | | | |
| | | | report_type_1:time_ranks | 0.44, CI: 0.07 0.81 | 0.02011 | 0.06499 |
| | | | **time_ranks** | **-0.79, CI: -1.17 -0.41** | **0.00005** | **0.00043** |
| AUDITORY | 0.044 | < 0.00001 | | | | |
| BIZARRENESS | 0.427 | < 0.00001 | | | | |
| | | | **report_type_1:time_ranks** | **-0.74, CI: -1.17 -0.31** | **0.00073** | **0.00956** |
| BODY | 0.076 | < 0.00001 | | | | |
| LIMITATIONS | 0.109 | < 0.00001 | | | | |
| | | | **time_ranks** | **-1.09, CI: -1.52 -0.65** | **< 0.00001** | **0.00001** |
| THOUGHT | 0.348 | < 0.00001 | | | | |
| | | | **report_type_1:time_ranks** | **0.63, CI: 0.24 1.02** | **0.00157** | **0.00956** |
| | | | time_ranks | -0.96, CI: -1.44 -0.48 | 0.00009 | 0.00049 |
| MOVEMENTS | 0.077 | < 0.00001 | | | | |
| | | | report_type_1:time_ranks | -0.51, CI: -0.97 -0.06 | 0.02658 | 0.07087 |
| SETTINGS | 0.178 | < 0.00001 | | | | |
| | | | report_type_1:time_ranks | -0.45, CI: -0.83 -0.07 | 0.02031 | 0.06499 |
| SOCIAL | 0.129 | < 0.00001 | | | | |
| SPACE | 0.212 | < 0.00001 | | | | |
| | | | report_type_1:time_ranks | -0.41, CI: -0.78 -0.04 | 0.03169 | 0.07243 |
| TACTILE | 0.063 | < 0.00001 | | | | |
| TIME | 0.176 | < 0.00001 | | | | |
| | | | **report_type_1:time_ranks** | **0.69, CI: 0.26 1.12** | **0.00179** | **0.00956** |
| | | | time_ranks | -0.87, CI: -1.31 -0.42 | 0.00015 | 0.00060 |
| VALENCE | 0.064 | < 0.00001 | | | | |
| | | | **time_ranks** | **0.59, CI: 0.27 0.91** | **0.00025** | **0.00079** |
| VISUAL | 0.190 | < 0.00001 | | | | |

***Table S11. Performance measures of the GLME model for the prediction of semantic dimensions across vigilance states in the discovery dataset, based on time.*** *The GLME model included vigilance state (report_type) as a regressor of interest, along with its interaction with time, and age, sex, education level, and the BADA score as covariates; q<0.05, False discovery Rate -FDR- correction. Significant effects after FDR correction are reported in bold.*

Table S12

| | Domain | Full Model adj $R^2$ | p-value Full model | coefficient name | coefficient beta | p-value coefficient | q-value coefficient |
|---|---|---|---|---|---|---|---|
| | DOMAIN2 | 0.066 | < 0.00001 | | | | |
| | DOMAIN4 | 0.023 | < 0.00001 | | | | |
| | Food | 0.009 | < 0.00001 | | | | |
| | | | | time_ranks | -0.63, CI: -1.08 -0.19 | 0.00539 | 0.07280 |
| | Objects | 0.049 | < 0.00001 | | | | |
| | | | | report_type_1:time_ranks | 0.84, CI: 0.01 1.67 | 0.04712 | 0.50258 |
| | | | | time_ranks | -0.96, CI: -1.73 -0.20 | 0.01344 | 0.08601 |
| | Reactions | 0.014 | 0.57757 | | | | |
| | DOMAIN12 | 0.015 | < 0.00001 | | | | |
| | Matter | 0.007 | 0.00011 | | | | |
| | Thriller | 0.029 | < 0.00001 | | | | |
| | Healthcare | 0.055 | < 0.00001 | | | | |
| | Locations | 0.061 | < 0.00001 | | | | |
| | Geometry | 0.051 | < 0.00001 | | | | |
| | Nature | 0.034 | < 0.00001 | | | | |
| | Fantasy | 0.001 | 0.15647 | | | | |
| | Measurements | 0.007 | 0.00004 | | | | |
| | Architecture | 0.032 | < 0.00001 | | | | |
| | Animals | 0.071 | < 0.00001 | | | | |
| | | | | time_ranks | -0.76, CI: -1.46 -0.05 | 0.03499 | 0.18664 |
| | Concerns | 0.035 | < 0.00001 | | | | |
| | Appearance | 0.074 | < 0.00001 | | | | |
| | Jobs | 0.049 | < 0.00001 | | | | |
| | | | | report_type_1:time_ranks | -0.62, CI: -1.21 -0.04 | 0.03732 | 0.50258 |
| | | | | time_ranks | 0.64, CI: 0.14 1.14 | 0.01232 | 0.08601 |
| | Drama | 0.072 | < 0.00001 | | | | |
| | Society | 0.020 | < 0.00001 | | | | |
| | | | | **time_ranks** | **-0.95, CI: -1.43 -0.46** | **0.00012** | **0.00384** |
| | Humanities | 0.098 | < 0.00001 | | | | |
| | | | | time_ranks | 0.67, CI: 0.18 1.16 | 0.00683 | 0.07280 |
| | Transportation | 0.019 | < 0.00001 | | | | |
| | Timing | 0.170 | < 0.00001 | | | | |
| | | | | report_type_1:time_ranks | 0.60, CI: 0.08 1.12 | 0.02261 | 0.50258 |
| | Technology | 0.031 | < 0.00001 | | | | |
| | Education | 0.047 | < 0.00001 | | | | |
| | Communication | 0.031 | < 0.00001 | | | | |
| | Slang | 0.015 | < 0.00001 | | | | |

***Table S12. Performance measures of the GLME model for the prediction of lexical domains across vigilance states in the discovery dataset, based on time.*** *The GLME model included vigilance state (report_type) as a regressor of interest, along with its interaction with time, and age, sex, education level, and the BADA score as covariates; q<0.05, False discovery Rate -FDR- correction. Significant effects after FDR correction are reported in bold.*

Table S13

| Anonymization code | Selected Words | Notes |
|---|---|---|
| NAME | Andrea | The names are listed in order: three male followed by three female names most commonly used in the year 2000 in the Italian Civil Registry, according to data from the Italian National Institute of Statistics (ISTAT, 2023. Indicatori demografici. *Istituto Nazionale di Statistica*). |
| | Francesco | |
| | Matteo | |
| | Alessia | |
| | Chiara | |
| | Martina | |
| PLACE | Torino | The cities are listed in geographic order: three from northern Italy, three from central Italy, and four from southern Italy, including the islands. |
| | Milano | |
| | Trieste | |
| | Firenze | |
| | Perugia | |
| | Roma | |
| | Napoli | |
| | Potenza | |
| | Bari | |
| | Palermo | |
| | Cagliari | |
| INSTITUTION | *fattoria* (farm) | Nouns referring to organization, institutes or services. The nouns are grouped and ordered by economic sector: two from the primary sector, six from the secondary, and six from the tertiary. |
| | *porto* (harbor) | |
| | *azienda* (company) | |
| | *cantiere* (construction site) | |
| | *bottega* (workshop) | |
| | *negozio* (store) | |
| | *ospedale* (hospital) | |
| | *banca* (bank) | |
| | *scuola* (school) | |
| | *università* (university) | |
| | *museo* (museum) | |
| | *teatro* (theater) | |
| | *biblioteca* (library) | |
| | *cinema* (cinema) | |

***Table S13. List of representative words chosen for the replacement of anonymization codes.*** *The first column lists the codes used to mask sensitive information in verbal reports, referring respectively to individuals, locations, and organizations. The second column presents the representative words selected to approximate the lexical information lost during anonymization. The third column provides additional information about the selected words.*

Table S14

| Variable Name | Internal Name | Variable Use | Variable Type | Variable level | Variable Description |
|---|---|---|---|---|---|
| AGENTIVITY | ACT | dependent | Likert 1-9, 1: extremely passive, 5: observer, 9: extremely active | report | semantic dimension |
| AROUSAL | ARO | dependent | Likert 1-9 | report | semantic dimension |
| AUDITORY | AUD | dependent | Likert 1-9 | report | semantic dimension |
| BIZARRENESS | BIZ | dependent | Likert 1-9 | report | semantic dimension |
| BODY | BOD | dependent | Likert 1-9 | report | semantic dimension |
| INCORPORATION | INC | dependent | Likert 1-9 | report | semantic dimension |
| LIMITATIONS | LIM | dependent | Likert 1-9 | report | semantic dimension |
| THOUGHT | THO | dependent | Likert 1-9 | report | semantic dimension |
| MOVEMENTS | MOV | dependent | Likert 1-9 | report | semantic dimension |
| SETTINGS | SET | dependent | Likert 1-9 | report | semantic dimension |
| SOCIAL | SOC | dependent | Likert 1-9 | report | semantic dimension |
| SPACE | SPA | dependent | Likert 1-9 | report | semantic dimension |
| TACTILE | TAC | dependent | Likert 1-9 | report | semantic dimension |
| TIME | TIM | dependent | Likert 1-9 | report | semantic dimension |
| VALENCE | VAL | dependent | Likert 1-9, 1: extremely negative, 5: neutral, 9: extremely positive | report | semantic dimension |
| VISUAL | VIS | dependent | Likert 1-9 | report | semantic dimension |
| DOMAIN2 | DOMAIN2 | dependent | Binary 0/1 | report | lexical domain |
| DOMAIN3 | DOMAIN3 | dependent | Binary 0/1 | report | lexical domain |
| DOMAIN4 | DOMAIN4 | dependent | Binary 0/1 | report | lexical domain |
| Food | DOMAIN6 | dependent | Binary 0/1 | report | lexical domain |
| Objects | DOMAIN9 | dependent | Binary 0/1 | report | lexical domain |
| Reactions | DOMAIN10 | dependent | Binary 0/1 | report | lexical domain |
| DOMAIN12 | DOMAIN12 | dependent | Binary 0/1 | report | lexical domain |
| Matter | DOMAIN14 | dependent | Binary 0/1 | report | lexical domain |
| Toponyms | DOMAIN15 | dependent | Binary 0/1 | report | lexical domain |
| Thriller | DOMAIN16 | dependent | Binary 0/1 | report | lexical domain |
| Healthcare | DOMAIN18 | dependent | Binary 0/1 | report | lexical domain |
| Locations | DOMAIN19 | dependent | Binary 0/1 | report | lexical domain |
| Geometry | DOMAIN20 | dependent | Binary 0/1 | report | lexical domain |
| Nature | DOMAIN21 | dependent | Binary 0/1 | report | lexical domain |
| Fantasy | DOMAIN27 | dependent | Binary 0/1 | report | lexical domain |
| Measurements | DOMAIN29 | dependent | Binary 0/1 | report | lexical domain |
| Architecture | DOMAIN32 | dependent | Binary 0/1 | report | lexical domain |
| Animals | DOMAIN33 | dependent | Binary 0/1 | report | lexical domain |
| Concerns | DOMAIN34 | dependent | Binary 0/1 | report | lexical domain |
| Appearance | DOMAIN35 | dependent | Binary 0/1 | report | lexical domain |
| Jobs | DOMAIN36 | dependent | Binary 0/1 | report | lexical domain |
| Conflicts | DOMAIN38 | dependent | Binary 0/1 | report | lexical domain |
| Drama | DOMAIN39 | dependent | Binary 0/1 | report | lexical domain |
| Society | DOMAIN41 | dependent | Binary 0/1 | report | lexical domain |
| Humanities | DOMAIN42 | dependent | Binary 0/1 | report | lexical domain |
| Transportation | DOMAIN44 | dependent | Binary 0/1 | report | lexical domain |
| Timing | DOMAIN45 | dependent | Binary 0/1 | report | lexical domain |
| Technology | DOMAIN46 | dependent | Binary 0/1 | report | lexical domain |

| | | | | | |
|---|---|---|---|---|---|
| Education | DOMAIN47 | dependent | Binary 0/1 | report | lexical domain |
| Transaction | DOMAIN48 | dependent | Binary 0/1 | report | lexical domain |
| Communication | DOMAIN49 | dependent | Binary 0/1 | report | lexical domain |
| Slang | DOMAIN50 | dependent | Binary 0/1 | report | lexical domain |
| Discovery dataset | dataset1 | N/A | N/A | N/A | Name of the main dataset, collected from 03/2020 to 04/2024 |
| Test dataset | dataset2 | N/A | N/A | N/A | Name of the test dataset, collected during lockdown (04/2020) |
| Vigilance States | report_type | independent | Categorical, 0 for wakefulness reports, 1 for dreams | report | encode in dataset1 the difference between dream and wakefulness reports |
| Experiment | Experiment | independent | Categorical, 0 for the main dataset, 1 for the lockdown dataset | report | main dataset (dataset1) is encoded with '0', lockdown dataset (dataset2) with '1' |
| Participant | Subj | independent | categorical | subject | ID of each participant, used to model random effects |
| Time | time_ranks | independent | continuous, range 0..1 | report | time of acquisition of reports in dataset 1 since beginning, converted in ranks and normalized to 1 |
| Sex | Sex | independent | Categorical, 0 for female participants, 1 for male participants | subject | biological sex assessed by self-report |
| Age | Age | independent | continuous, in yrs | subject | chronological age |
| Education | Education | independent | continuous, in yrs | subject | highest degree obtained |
| Trait anxiety levels | STAI | independent | continuous | subject | C. D. Spielberger et al., Rev. Interam. Psicol. 5 (1971). |
| Perceived sleep quality | PSQI | independent | continuous | subject | D. J. Buysse et al., Psychiatry Res. 28, 193–213 (1989). |
| Attitude towards dreaming | ATD | independent | continuous | subject | K. Bulkeley, M. Schredl, Int. J. Dream Res. 12, 7 (2019). |
| Mind-Wandering (spontaneous, deliberate) | MW | independent | continuous | subject | K. Christoff et al., Nat. Rev. Neurosci. 17, 718–731 (2016). |
| Verbal memory | BSRT | independent | continuous | subject | H. Babcock, L. Levy, Test and Manual of Directions; the Revised Examination for the Measurement of Efficiency of Mental Functioning (Stoelting, Wood Dale, IL, US, 1940). |
| Visuospatial memory | ROCFr | independent | continuous | subject | A. Rey, Arch. Psychol. (Geneve) 28, 215–285 (1941). |
| Subjective circadian preference | MEQ | independent | continuous | subject | J. A. Horne, O. Ostberg, Int. J. Chronobiol. 4, 97–110 (1976). |
| Vividness of visual imagery | VVIQ | independent | continuous | subject | D. F. Marks, Br. J. Psychol. 64, 17–24 (1973). |
| Vulnerability to cognitive interference | SCWT | independent | continuous | subject | F. Scarpina, S. Tagini, Front. Psychol. 8, 557 (2017). |
| BADA | WC_BADA | independent | continuous | subject | Log10 count of words used in the BADA test |
| Word Count per participant | WC_subj | independent | continuous | subject | average Log10 count of words across reports of a given participant. It was used when comparing dataset2 with dataset1. |
| PC#1 from actigraphy | PC1_report | independent | continuous | report | sleep fragmentation - PC1 scores of actigraphic data of the night prior the report |
| PC#2 from actigraphy | PC2_report | independent | continuous | report | long light sleep - PC2 scores of actigraphic data of the night prior the report |
| PC#3 from actigraphy | PC3_report | independent | continuous | report | stable advanced sleep - PC3 scores of actigraphic data of the night prior the report |
| PC#4 from actigraphy | PC4_report | independent | continuous | report | unstable advanced sleep - PC4 scores of actigraphic data of the night prior the report |

***Table S14. Comprehensive description and naming conventions of all collected variables.*** *Overview of all collected variables and their naming conventions. The first column lists variable names as referenced in the main text; the second column shows their corresponding internal names used in the repository. Columns 3 to 5 provide information on variable type, scale, and source. The final column includes detailed descriptions and, where applicable, references.*
